# Chronic Ethanol Drinking Alters Medial Prefrontal Cortex and Nucleus Accumbens Astrocyte Translatome and Extracellular Matrix Glycosaminoglycans

**DOI:** 10.1101/2025.09.30.679577

**Authors:** Joel G. Hashimoto, Angela R. Ozburn, Cheryl Reed, Jason Erk, Yuefan Song, Jiyuan Yang, Ke Xia, Fuming Zhang, Yun Yu, Suzanne S. Fei, Lina Gao, Robert J. Linhardt, Tamara J. Phillips, Marina Guizzetti

## Abstract

Alcohol Use Disorder is a leading preventable cause of morbidity and mortality, yet knowledge of mechanisms driving ethanol-related neuroplasticity remains incomplete. While research has traditionally focused on neuronal signaling, emerging evidence implicates astrocytes in addiction-related adaptations. Here, we investigated the astrocyte-specific molecular consequences of chronic ethanol consumption in the prefrontal cortex and nucleus accumbens, two brain regions critical for executive control and reward processing. Using Translating Ribosome Affinity Purification RNA-seq and bulk RNA-seq in Aldh1l1-EGFP/Rpl10a mice, expressing an EGFP tag on astrocyte ribosomes, we identified hundreds of differentially translated astrocytic genes following chronic continuous two-bottle choice ethanol drinking. Sex-specific analyses revealed greater astrocytic changes in the female PFC and male NAc. Pathway enrichment highlighted extracellular matrix remodeling, synaptic signaling, mitochondrial function, and immune-related pathways. Analyses of individual drinking levels further demonstrated distinct correlations between ethanol intake and astrocytic translation. The major components of the brain extracellular matrix are chondroitin sulfate proteoglycans, produced primarily by astrocytes and covalently bound to chondroitin sulfate glycosaminoglycan chains. Complementary mass spectrometry/liquid chromatography analyses of chondroitin sulfate, heparan sulfate, and hyaluronic acid glycosaminoglycan disaccharides revealed ethanol-induced alterations in chondroitin sulfate glycosaminoglycan sulfation patterns, with additional baseline differences identified between selectively bred high– and low-ethanol preference lines. Together, these findings indicate that astrocytes undergo profound sex– and region-specific adaptations to chronic ethanol, implicating extracellular matrix and glycosaminoglycan remodeling as key risk-factors for and mediators of chronic ethanol-related neuroplasticity.

## 1. Introduction

Alcohol Use Disorder (AUD) is a major public health problem that imposes a significant burden on individuals, families, and society. In the United States, excessive alcohol use accounts for more than 178,000 deaths annually, making alcohol one of the leading preventable causes of death (ARDI Alcohol-Attributable Deaths, US | CDC, n.d.). Although alcohol adversely affects multiple organ systems, AUD is primarily classified as a psychiatric disorder. With continued drinking, progressive changes occur in brain structure and function (Koob and Volkow, 2016). Several brain regions have been implicated in different aspects of alcohol drinking and AUD (Koob and Volkow, 2016). These alterations compromise brain function and drive the transition from controlled, occasional use to chronic misuse, which can become difficult to regulate and ultimately lead to AUD (George and Koob, 2017). The persistence of these changes suggests stable behavioral adaptations and neuroplasticity mediated through transcriptional mechanisms.

Notably, the consequences of alcohol (ethanol; EtOH) on brain function have long remained focused on neurotransmitter systems and neuronal cell signaling. Transcriptional profiles of brain tissues from post-mortem studies (AUD vs control) and animal models have revealed a number of important EtOH-associated genes for mechanistic study (e.g. Farris et al., 2015; Colville et al., 2017, 2018; Ferguson et al., 2018; Pozhidayeva et al., 2020; Walter et al., 2021; Anderson et al., 2025; Gottlieb et al., 2025). Recent transcriptome findings indicate that glial cells, such as astrocytes, microglia, and oligodendrocytes, which have distinct and important roles in brain function and behavior, also play an important role in drinking (Erickson et al., 2019, 2021; Brenner et al., 2020; Warden et al., 2020, 2025; van den Oord et al., 2023). In the brain, astrocytes are the most abundant glial cells, representing 20-40% of the total number of cells. Astrocytes play a central role in synaptic regulation, metabolic support, and extracellular matrix (ECM) organization (Herculano-Houzel, 2014; Chung et al., 2024). Emerging research indicates that astrocytes contribute to the pathophysiology of alcohol and drug addiction (Kruyer et al., 2023; Holt and Nestler, 2024; Guizzetti et al., 2025). Chemogenetic manipulation of astrocytes in the prefrontal cortex (PFC) and nucleus accumbens (NAc) is sufficient to alter EtOH drinking (Bull et al., 2014; Erickson et al., 2021). However, the extent to which astrocytes adapt to chronic EtOH drinking remains poorly understood.

The roles of astrocytes as key mediators of brain development, function, and plasticity started emerging in the early 1990s. Since then, the known physiological functions of astrocytes have been steadily growing. Astrocytes ensheathe most synapses and regulate neurotransmitter levels, as they express high-affinity neurotransmitter transporters and contain enzymes needed for neurotransmitter synthesis and metabolism. Perisynaptic astrocyte processes express several metabotropic and ionotropic neurotransmitter receptors (Agulhon et al., 2008). Astrocytes are also involved in synapse development and remodeling through the release of several factors (Allen and Eroglu, 2017). Astrocytes surround brain blood vessels with their end-feet and regulate local blood flow and the maintenance of the blood brain barrier (MacVicar and Newman, 2015; Cheslow and Alvarez, 2016; Mishra, 2017), and have transporters and channels for ions and water (Kimelberg, 2010; Schousboe et al., 2013).

Glycosaminoglycans (GAGs) are long, unbranched polysaccharides consisting of repeating disaccharide units present on the cell surface and in the ECM (Thompson et al., 2017). Three forms of GAGs relevant to neuronal plasticity are chondroitin sulfate (CS-GAGs), heparan sulfate (HS-GAGs), and hyaluronic acid (HA). Chondroitin sulfate proteoglycans (CSPGs) of the lectican family (neurocan, brevican, versican, and aggrecan) are the most abundant proteins of the central nervous system ECM. Brevican and neurocan are brain-specific CSPGs, the most abundant lecticans in the brain, and primarily expressed and released by astrocytes (Zhang et al., 2021). Lecticans interact with numerous factors in the extracellular space (Bartus et al., 2012) and are major components of perineuronal nets (PNNs). Lecticans consist of core-proteins attached to one or more linear chondroitin sulfate glycosaminoglycan (CS-GAG) chains formed by repeated glucuronic acid and N-acetylgalactosamine disaccharides modified by sulfation (Prydz and Dalen, 2000). CS-GAGs, and in particular C4S-GAGs are inhibitors of axonal growth and regeneration, while the digestion of CS-GAGs by the bacterial enzyme chondroitinase ABC promotes recovery after brain and spinal cord injury in animal models (Carulli et al., 2005; Zhao and Fawcett, 2013). Heparan sulfate proteoglycans (HSPGs) can be membrane-bound or secreted and consist of a core protein covalently attached to long linear HS-GAG chains (Sarrazin et al., 2011). HSPGs are involved in a wide range of cellular processes by direct interactions with different binding partners. Most of these interactions occur in a HS-dependent and specific manner (Xu and Esko, 2014). HSPGs are expressed in a brain region– and cell type-specific manner and are important players in the formation and function of synaptic connections (Condomitti and de Wit, 2018). HS-GAGs are composed of repeating disaccharide units of N-acetylglucosamine and glucuronic acid that can be modified by N-sulfation and three sites of O-sulfation (Yu et al., 2017). Hyaluronic acid (HA) is a large, unbranched, non-sulfated GAG formed by the repeating disaccharide unit N-acetylglucosamine and N-glucuronic acid. HA is the only GAG that is not covalently bound to a core protein and serves as a backbone for lecticans, all of which have HA binding sites. HA also has binding sites for membrane receptors and is a major component of the PNNs (Miyata and Kitagawa, 2017).

Many functions of astrocytes have been identified using *in vitro* astrocyte cultures (Pfrieger and Barres, 1997; Ullian et al., 2001; Christopherson et al., 2005; Allen et al., 2012) because the study of astrocytes *in vivo* has been limited by the lack of appropriate molecular and genetic tools. However, new genetic tools allow for the selective manipulation and imaging of astrocytes and the isolation of *ex vivo* astrocyte RNA (Doyle et al., 2008; Srinivasan et al., 2016; Chai et al., 2017). Of particular note, the aldehyde dehydrogenase 1 family member L1 (*Aldh1l1*) gene was identified as a highly specific marker for astrocytes with a much broader pattern of astrocyte expression than the traditional marker *Gfap* (Cahoy et al., 2008). Here we use a transgenic mouse model expressing an EGFP-tagged Rpl10a ribosomal protein targeted to cells expressing *Aldh1l1* to evaluate gene expression specifically occurring in astrocytes (Doyle et al., 2008; Heiman et al., 2008a). This transgenic model has been used to explore astrocyte gene translation at different stages of development and aging, after alterations in sleep patterns, and after developmental EtOH exposure (Bellesi et al., 2015; Morel et al., 2017; Clarke et al., 2018; Hashimoto et al., 2025).

The PFC and NAc are critical hubs in executive function and reward processing, respectively, and are both important for AUD (Koob and Volkow, 2016). Despite their importance, astrocyte-specific translational responses to EtOH in these regions remain understudied. Furthermore, sex differences in EtOH intake and associated neuroadaptations are well-documented (Hitzemann et al., 2022), yet their astrocytic underpinnings are not well understood. Additionally, ECM remodeling and alterations in glycosaminoglycan (GAG) composition are increasingly recognized as contributors to neuroplasticity (Carulli et al., 2005; Dzyubenko and Hermann, 2023), but their role in EtOH-related astrocytic adaptations has not been comprehensively investigated.

In the present study, we carried out Translating Ribosome Affinity Purification (TRAP) RNA-seq and traditional RNA-seq to examine astrocyte-specific translational changes in the PFC and NAc following chronic EtOH intake. We also evaluated sex differences in these adaptations and assessed correlations with individual EtOH drinking levels. We then quantified HA, CS-, and HS-GAG composition of the PFC and NAc of TRAP and high-EtOH preference (HP) mice to explore ECM-related changes caused by long-term EtOH intake. Finally, to learn whether ECM constituents conferred risk for high EtOH drinking, we compared GAG composition in EtOH naïve high– and low-ethanol preference (HP/LP) mouse lines. We report that chronic EtOH consumption produces sex– and region-specific astrocytic changes, with enrichment in pathways related to ECM remodeling, synaptic signaling, and metabolic regulation, implicating ECM and GAG remodeling as key risk-factors for and mediators of chronic EtOH-related neuroplasticity.

## 2. Methods

### 2.1 Animals

All experiments involved experimentally naïve adult male and female mice. Methods were consistent with guidelines from the National Institutes of Health and approved by the Institutional Animal Care and Use Committee of the VA Portland Health Care System (VAPORHCS).

#### Aldh1l1-EGFP/Rpl10a Mice

Adult hemizygous Aldh1l1-EGFP/Rpl10a mice and their wild-type conspecifics were purchased from The Jackson Laboratory (Strain # 030247) and housed within the VAPORHCS animal facility. Aldh1l1-EGFP-Rpl10a transgenic mice were used because they express a modified ribosomal protein with an enhanced green fluorescent protein (EGFP) tag only in cells expressing the astrocytic marker *Aldh1l1*, allowing the harvest and quantitative analysis of astrocyte-specific translating RNA.

The mice were allowed to acclimate to the vivarium for one week prior to being set up as hemizygous x wild-type breeder pairs. At weaning, all offspring were ear-punched for individual tracking and genotyping. Genotyping was performed to identify each mouse as wild-type or hemizygous transgenic. All genotyping was performed using traditional PCR methods as described by Doyle et al. (2008), with primers suggested by The Jackson Laboratory.

#### High and Low EtOH Preference Mice

Selectively bred high EtOH preference (HP) and low EtOH preference (LP) mice were created by the Animal & Resource Core of the Portland Alcohol Research Center (Colville et al., 2017 and 2018; Anderson et al., 2025). Methods and outcomes for the selection are detailed in Anderson et al. (2025). The selection phenotype was average EtOH preference compared to water from the second and fourth offering of 10% EtOH (mice were offered 5% EtOH vs. water for 4 days and then 10% EtOH vs. water for 4 days). HP and LP mice used in these studies were from selection generation 5 (S5), from the second replication of these lines.

### 2.2 Chronic EtOH Intake

A total of 205 mice were tested in this procedure. Mice were an average of 63 days old at the start of the drinking study (range 48-71). Animals were placed on a 12:12 h light:dark cycle with lights on at 0600 h and off at 1800 h. A schematic of the experimental timeline is shown in Figure 1A. A 2-bottle choice (2-BC), 24-h continuous access drinking protocol was used with consecutive steps of habituation to single housing and acclimation to drinking from two water-filled 25-ml graduated cylinders fitted with sipper tubes for 4 days, and measurement of preference for water vs. EtOH in water (v/v) for 4 days each. On test days 5-8, mice were offered water vs. 5% EtOH in water (v/v) to acclimate them to EtOH drinking at a lower concentration (test days 5-8) and then offered water vs. 10% EtOH from test day 9 through 92 or 93. The left-or right-side position of the EtOH tube relative to the water tube was alternated every other day to account for potential drinking side preference. Animals were weighed and tubes containing fresh solutions were placed on the cages every 4 days. Mice had *ad libitum* access to standard rodent chow throughout the testing period. Unoccupied control cages were handled identically to test cages to correct for leakage and evaporation. The amount of fluid in each tube (ml) was recorded daily and calculated from tube volumes was daily EtOH consumed (g/kg), total volume consumed (water + EtOH; ml), and preference for EtOH (preference ratio: ml of EtOH consumed divided by total volume consumed). The total number of EtOH-drinking animals tested was 130: 24 HP mice (12 per sex), 0 LP mice, and 106 Aldh1l1-EGFP/Rpl10a (54 males, 52 females) mice. Additionally, a control group received access to two bottles of water over the entire course of testing. The total number of control animals tested was 75: 12 HP (6 per sex), 12 LP (6 per sex), and 51 Aldh1l1-EGFP/Rpl10a (25 males, 26 females) mice. This study was focused on the impact of high EtOH intake, compared to water only, on molecular signature; thus, the low intake LP line was not tested for water vs. EtOH drinking. However, LP and HP lines in the water group were compared for innate differences between the lines.

**Figure 1.**
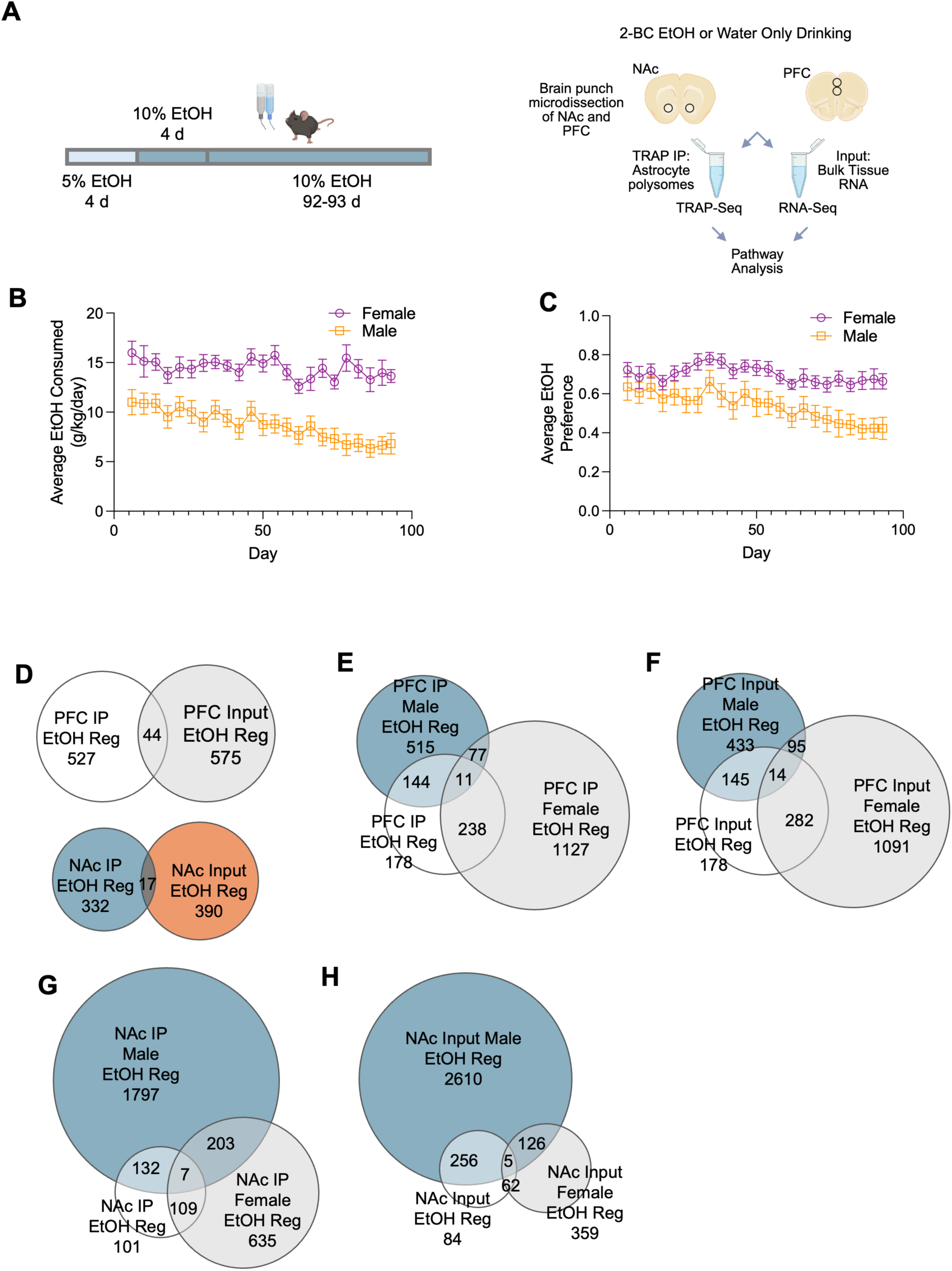
Long-term EtOH drinking in Aldh1l1-EGFP/Rpl10a mice alters PFC and NAc astrocyte gene translation. **A**. Left panel: Aldh1l1-EGFP/Rpl10a mice underwent 2-BC, 24 h unlimited access drinking of 10% EtOH for 92-93 days. An additional set of animals had access to water in both bottles throughout the study to allow comparisons of 2-BC EtOH drinking and ethanol naïve mice. Right panel: Snap frozen brains from 2-BC EtOH or water only drinking animals underwent brain punch microdissection for NAc and PFC. From each tissue, astrocyte translating RNA was immunoprecipitated (IP) by the TRAP procedure. In addition, RNA from the brain homogenate (Input) was collected to determine the bulk-tissue expression profile. Created in BioRender.com. **B**. Daily ethanol consumption was recorded; the data points shown in the figure are the average of 4 days of drinking during the 2-BC 10% ethanol drinking days in male and female TRAP mice. **C**. Each data point represents the average of 4 days for EtOH preference during 10% EtOH drinking. **D**. RNA-Seq on IP and Input fractions identified astrocyte differentially translated RNA as well as bulk-tissue differentially expressed RNA. Input and IP fractions showed little overlap in the PFC and NAc.For each of the Venn diagrams, the number of DT or DE genes in each comparison are listed. **E**. TRAP-seq results in the PFC showed more DT genes in females compared to males. **F**. Similarly, input RNA-seq results in the PFC also showed more DE genes in females compared to males. **G**. In the NAc, males showed higher number of DT genes compared to females. **H**. In NAc input RNA-seq results, males also showed higher number of DE genes than females.

### 2.3 Tissue Collection

Mice were randomly assigned to two dissection groups, each containing equal numbers of mice of each genotype, sex and treatment (i.e., water only or water vs. EtOH). Immediately after the final drinking measurement, on day 92 for half of the mice and day 93 for the other half, whole brains were collected, snap frozen in isopentane cooled in a dry ice isopropanol slurry and stored at –80° C.

### 2.4 Translating-RNA Affinity Purification (TRAP)

Brain punch microdissections were carried out on 300 µm coronal sections using a 1.2 mm diameter punch tool and stored at –80° C until processing. TRAP immunoprecipitation (IP) was carried out on tissue punches as previously described (Doyle et al., 2008; Heiman et al., 2008b; Hashimoto et al., 2025).

### 2.5 RNA-Seq Data Processing and Analysis

RNA-seq libraries were generated using the Illumina Stranded mRNA Prep kit, profiled on a 4200 Tapestation (Agilent) and quantified by real time PCR using a commercial kit (Kapa Biosystems / Roche) on a StepOnePlus Real Time PCR Workstation (ABI/Thermo). Libraries were then sequenced on a NovaSeq 6000 (Illumina). Fastq files were assembled from the base call files using bcl2fastq (Illumina). Library preparation and RNA sequencing were carried out by the Integrated Genomics Laboratory at OHSU.

Differential expression analysis was performed by the ONPRC Bioinformatics & Biostatistics Core. The quality of the raw sequencing files was evaluated using FastQC (Andrews, 2010) combined with MultiQC (Ewels et al., 2016) (http://multiqc.info/). Trimmomatic (Bolger et al., 2014) was used to remove any remaining Illumina adapters. Reads were aligned to Ensembl’s Mus musculus GRCm39 genome along with its corresponding annotation, release 110. The program STAR (Dobin et al., 2013) (v2.7.10b_alpha_220111) was used to align the reads to the genome. STAR has been found to perform well compared to other RNA-seq aligners (Engström et al., 2013). Since STAR utilizes the gene annotation file, it also calculated the number of reads aligned to each gene. RNA-SeQC and another round of MultiQC were utilized to ensure alignments were of sufficient quality.

Gene-level differential expression analysis was performed in open-source software R (Version 4.3.2). Gene-level raw counts were filtered to remove genes with extremely low counts in many samples following the published guidelines (Chen et al., 2016), normalized using the trimmed mean of M-values method (TMM) (Robinson and Oshlack, 2010), and transformed to log-counts per million with associated observational precision weights using the voom method (Law et al., 2014). Gene-wise linear models with primary variables EtOH consumption and sex, adjusting for trap procedures, were employed for differential expression analyses using limma with empirical Bayes moderation (Ritchie et al., 2015) and false discovery rate (FDR) adjustment (Benjamini and Hochberg, 1995). Differential expression data were analyzed using Ingenuity Pathway Analysis software (QIAGEN Inc.), using a cutoff for significant regulated genes of p value < 0.05.

### 2.6 Glycosaminoglycan Quantification

Unsaturated disaccharide standards of CS, HS, and HA were purchased from Iduron (UK). Actinase E was obtained from Kaken Biochemicals (Japan). Chondroitin lyase ABC from Proteus vulgaris was expressed in the laboratory of Fuming Zhang. Recombinant Flavobacterial heparin lyases I, II, and III were also expressed in our laboratory using Escherichia coli strains provided by Jian Liu (College of Pharmacy, University of North Carolina). 2-Aminoacridone (AMAC), sodium cyanoborohydride were obtained from Sigma-Aldrich (St. Louis, MO, USA). All solvents were HPLC grade.

Frozen brain samples were freeze dried and proteolyzed at 55°C with 200 µL of 10 mg/mL actinase E for 24 h, followed by actinase E deactivation at 100°C for 30 min. The digested solution was transferred to a 10 kDa molecular weight cut off (MWCO) spin tube. CHAPS was added (final concentration 2%) and mixed with the digested solution in the filter of the spin tube. The filter unit was washed three times with 400 μL distilled water before adding 300 µL of digestion buffer (50 mM ammonium acetate containing 2 mM calcium chloride adjusted to pH 7.0). Recombinant heparin lyaseI, II, III (pH optima 7.0−7.5) and recombinant chondroitin lyase ABC (pH optimum 7.4, 10 mU each) were added to each filter unit containing sample and mixed well. All samples were incubated at 37° C for 24 h. The enzymatic digestion was terminated by ultrafiltration through the 10 kDa spin tube. The filtrate was collected, the filter unit was washed twice with 200 μL distilled water, and all the filtrates containing the disaccharide products were combined and freeze dried.

Freeze dried samples were AMAC-labeled by adding 10μL of 0.1 M AMAC in DMSO/acetic acid (17/3, v/v) incubating at room temperature for 10 min, followed by adding 10μL of 1 M aqueous sodium cyanoborohydride and incubating for 1 h at 45°C. A mixture containing all 17-disaccharide standards prepared at 0.5ng/μL was similarly AMAC-labeled and used for each run as an external standard. After the AMAC-labeling reaction, the samples were centrifuged, and each supernatant recovered.

Liquid chromatography was performed on an Agilent 1200 LC system at 45 °C using an Agilent Poroshell 120 ECC18 (2.7 μm, 3.0 × 50 mm) column. Mobile phase A (MPA) was 50 mM ammonium acetate aqueous solution, and the mobile phase B (MPB) was methanol. The mobile phase passed through the column at a flow rate of 300 μL/min. The gradient was 0-10 min, 5-45% B; 10-10.2 min, 45-100% B; 10.2-14min, 100% B; 14-22 min, 100-5% B. Injection volume was 5 μL.

A triple quadrupole mass spectrometry system equipped with an ESI source (Thermo Fisher Scientific, San Jose, CA) was used as the detector. The online MS analysis was at the Multiple Reaction Monitoring (MRM) mode. MS parameters: negative ionization mode with a spray voltage of 3000 V, a vaporizer temperature of 400°C, and a capillary temperature of 250°C. CS and HS GAG disaccharides are presented as percent of total CS or HS and relative proportions of CS, HS, and HA as previously described (Zhang et al., 2021; Hashimoto et al., 2023, 2025).

### 2.7 Statistical Analysis

GAG proportions and 2-BC daily EtOH drinking parameters (volume, EtOH g/kg/day, and preference ratio), were analyzed using linear intercept-only mixed-effects models using the lme4 package in R (Bates et al., 2015; Goeke et al., 2018). For EtOH drinking parameters, ‘day’ was included as a random effect in the mixed-effects model. Effects of sex, EtOH treatment, and sex by EtOH treatment interactions were assessed by p-values generated by chi-squared test with significance determined p < 0.05. For qRT-PCR confirmation of RNA-Seq data, two-way ANOVA with sex by treatment interactions or t-tests were used in PRISM 8.

## 3. Results

### 3.1 2-BC choice drinking

EtOH intake and preference data are shown in Figure 1 for Aldh1l1-EGFP/Rpl10a (astrocyte TRAP) mice that underwent a 24 h unlimited access 2-BC procedure with 10% EtOH vs. water for 92-93 days. Data points shown are for 4-day averages. Figure 1A (left side) shows the time course of behavioral testing. Linear mixed-effects modeling found significant sex differences in EtOH consumption (g/kg/day; χ^2^(1) = 746.6, p = 2.2×10^−164^; Figure 1B), EtOH preference ratio (χ^2^(1) = 180.9, p = 3.1×10^−41^; Figure 1C), and total volume consumed (mL/day; χ^2^(1) = 161.3, p = 5.8×10^−37^; data not shown). Female astrocyte TRAP mice drank on average 14.3 g/kg/day (SD = 3.93), while males drank 8.7 g/kg/day (SD = 3.97). EtOH preference in females was 0.69 (SD = 0.17) and in males 0.55 (SD = 0.25). Total volume consumed per day was 6.07 mL (SD = 1.31) in females and 5.66 mL (SD = 0.97) in males.

### 3.2 TRAP and RNA-Seq

To understand the effects of long-term EtOH drinking on astrocyte translation, TRAP was run on PFC and NAc punches to isolate astrocyte translating RNA and input (bulk tissue) RNA from 37 mice; 12 females and 13 males that underwent the 2-BC and 6 female and 6 male mice with water-only access. RNA-seq was run on a total of 148 samples: 2 brain regions (NAc and PFC) and two fractions (TRAP IP and input) from 37 mice. The average RNA integrity number (RIN) was 7.33 (SD: 1.34) with a range of 4.9-9.1 (Supplemental Table 1). A schematic of the experimental design is shown on the right side of Figure 1A.

### 3.3 2-BC EtOH drinking vs water drinking RNA-Seq and pathway analyses

Our initial analysis compared 2-BC and water-only controls for each brain region and fraction (Table 1). In the PFC, we identified 571 differentially translated (DT) genes in the astrocyte (IP) fraction (237 up-regulated by EtOH, 334 down-regulated) and 619 genes differentially expressed (DE) in the input fraction (242 up-regulated, 377 down-regulated; p < 0.05). In the NAc, we identified 349 DT genes in the IP fraction (161 up-regulated, 188 down-regulated) and 407 DE genes in the input fraction (168 up-regulated, 239 down-regulated; Table 1).

**Table 1.** Significant Gene Counts.

| Comparison | PFC<br>Ast. | PFC<br>Input | NAc<br>Ast. | NAc<br>Input |
| --- | --- | --- | --- | --- |
| EtOH Vs Water | 571 | 619 | 349 | 407 |
| EtOH Vs Water - Female | 1453 | 1482 | 954 | 552 |
| EtOH Vs Water - Male | 747 | 687 | 2139 | 2997 |
| Drinking - All Days | 643 | 419 | 1025 | 1639 |
| Drinking - All Days Female | 648 | 728 | 595 | 636 |
| Drinking - All Days Male | 844 | 716 | 1501 | 1629 |
| Drinking - First 25 Days | 815 | 394 | 814 | 754 |
| Drinking - First 25 Days Female | 498 | 551 | 621 | 576 |
| Drinking - First 25 Days Male | 889 | 561 | 1334 | 692 |
| Drinking - Last 25 Days | 739 | 506 | 1258 | 1656 |
| Drinking - Last 25 Days Female | 787 | 885 | 525 | 445 |
| Drinking - Last 25 Days Male | 1035 | 695 | 1737 | 2342 |

Number of significant genes in PFC astrocytes (PFC Ast.), PFC bulk tissue (PFC Input), NAc astrocytes (NAc Ast.), and NAc bulk tissue (NAc Input). The first three rows above the dashed line are categorical analysis comparing EtOH drinking to water drinking animals. Rows below the dashed line are the number of genes either positively or negatively correlated with EtOH drinking across all days, the first 25 days, or the last 25 days of drinking.

Comparison of DT and DE genes found very little overlap with only 17 genes in common in the NAc and 44 genes in common in the PFC (Figure 1D). When looking at the effects of 2-BC drinking within each sex subgroup, we identified more DT and DE genes than in the sexes combined analyses, with more DT and DE genes in the female PFC than male PFC and more DT and DE genes in the male NAc than female NAc (Table 1). Comparison of overlap between the different sexes within each brain region and fraction are presented in Figure 1E-H.

Significantly DT and DE genes from each brain region were used in Ingenuity Pathway Analysis (IPA, Qiagen) for combined as well as sex separated analyses. IPA canonical pathway enrichment analysis uses the direction of regulation to calculate an activation z-score for each enriched pathway which provides information on whether a given pathway is expected to be activated or inhibited. The top canonical pathways from IPA analysis of DT genes (astrocyte translating RNA) are shown in Figure 2 where the size of the dot corresponds to the enrichment ratio (regulated genes/total genes in pathway), the color corresponds to the magnitude of the p-value with brighter colors indicating higher significance, and location on the x-axis indicating expected activation/inhibition status. In the PFC, “Complement System” is the top enriched pathway, with “Dopamine Degradation” and “Integrin cell surface interactions” among the top enriched pathways (Figure 2A). When looking at females alone (Figure 2B), PFC DT genes are enriched in translation related pathways as well as “Glutaminergic Receptor Signaling” and “Calcium Transport I”. In males, the PFC related DT pathways were “Extracellular matrix organization” and “Integrin cell surface interactions” and enrichment of “Axonal Guidance Signaling” (Figure 2C).

**Figure 2.**
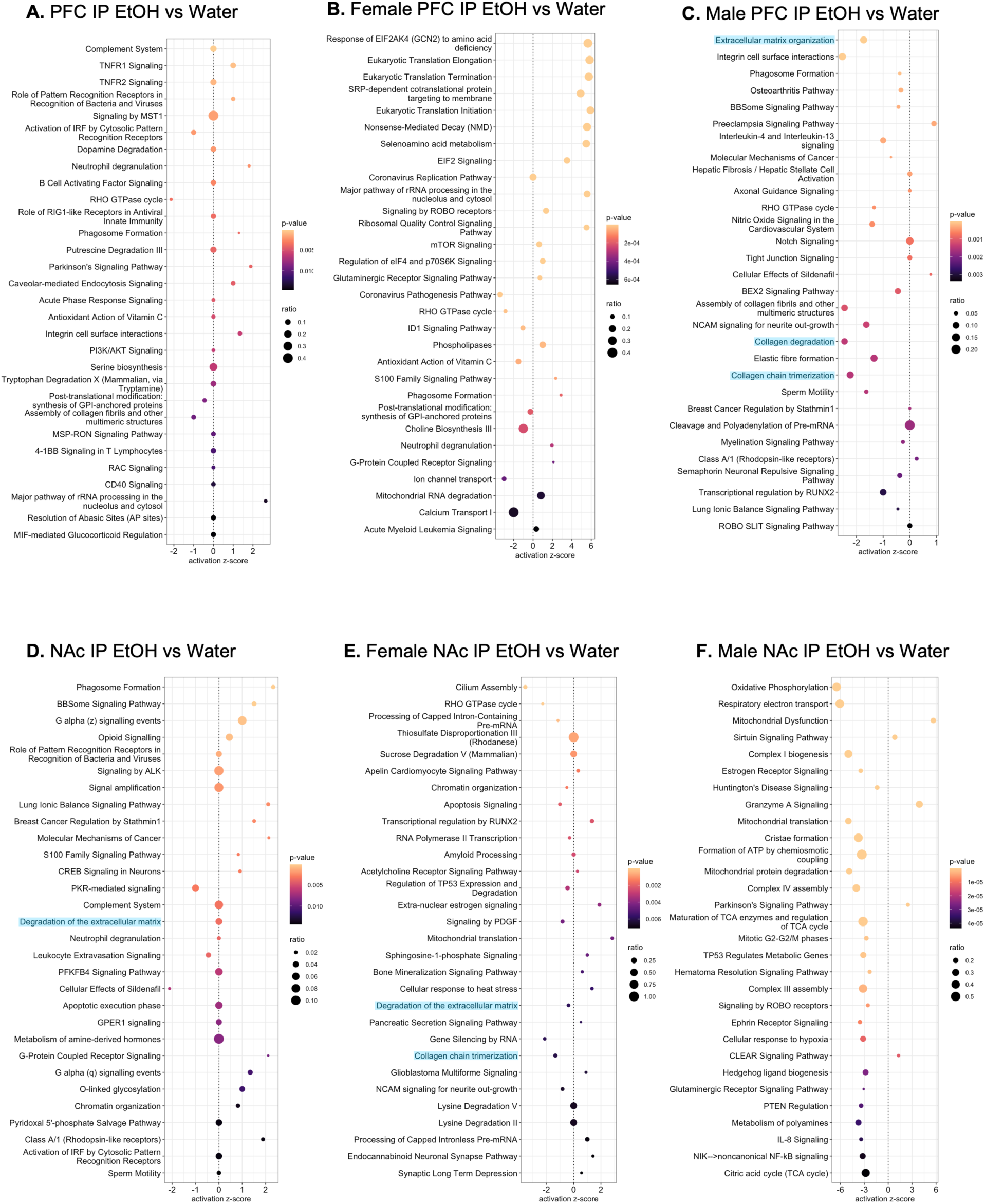
Top 30 Canonical Pathways enriched in astrocyte DT genes altered by EtOH drinking. DT genes were used for IPA assessment of canonical pathway enrichment and expression level was used to determine the activation z-score, inferring whether a pathway is up or down regulated by ethanol. **A**. Top enriched canonical pathways in DT genes in analysis of the PFC. **B**. Top enriched canonical pathways in DT genes in the PFC of females only. **C**. Top enriched canonical pathways in DT genes in the PFC of males only. **D**. Top enriched canonical pathways in DT genes in analysis of the NAc. **E**. Top enriched canonical pathways in DT genes in the NAc of females only. **F**. Top enriched canonical pathways in DT genes in the NAc of males only. Pathways related to the ECM are highlighted.

In the NAc, “Phagosome Formation” and “BBSome Signaling Pathway” were the top enriched categories and had positive activation z-scores (Figure 2D). BBSome is a protein complex involved in the biogenesis and homeostasis of the primary cilia; non-motile “antennae” that sense extracellular signals. Other enriched pathways of note are “Opioid Signaling”, “Complement System”, and “Degradation of the extracellular matrix”. In the NAc of females, 2-BC vs water DT genes showed evidence of decreased activation in “Cilium assembly” and “RHO GTPase cycle” pathways (Figure 2E). Previous bulk-tissue RNA-seq analysis of the central amygdala after 3-month 2-BC drinking identified primary cilia pathways correlated with EtOH drinking in females (Hitzemann et al., 2020, 2022). Other pathways of note include “Degradation of the extracellular matrix” and “NCAM signaling for neurite outgrowth”. In the NAc of males, top enriched categories were “Oxidative Phosphorylation”, “Respiratory electron transport”, and “Mitochondrial Dysfunction” (Figure 2F).

### 3.4 Pathway analysis correlating EtOH drinking levels and astrocyte functions

Figure 3A shows individual animal EtOH consumption averaged across all days during the 2-BC procedure (left panel), individual animal EtOH consumption averaged across the first 25 drinking days (middle panel), and individual animal EtOH consumption averaged across the last 25 days of EtOH drinking (right panel). Four-day averages for EtOH intake for these animals are shown in Figure 1B.

**Figure 3.**
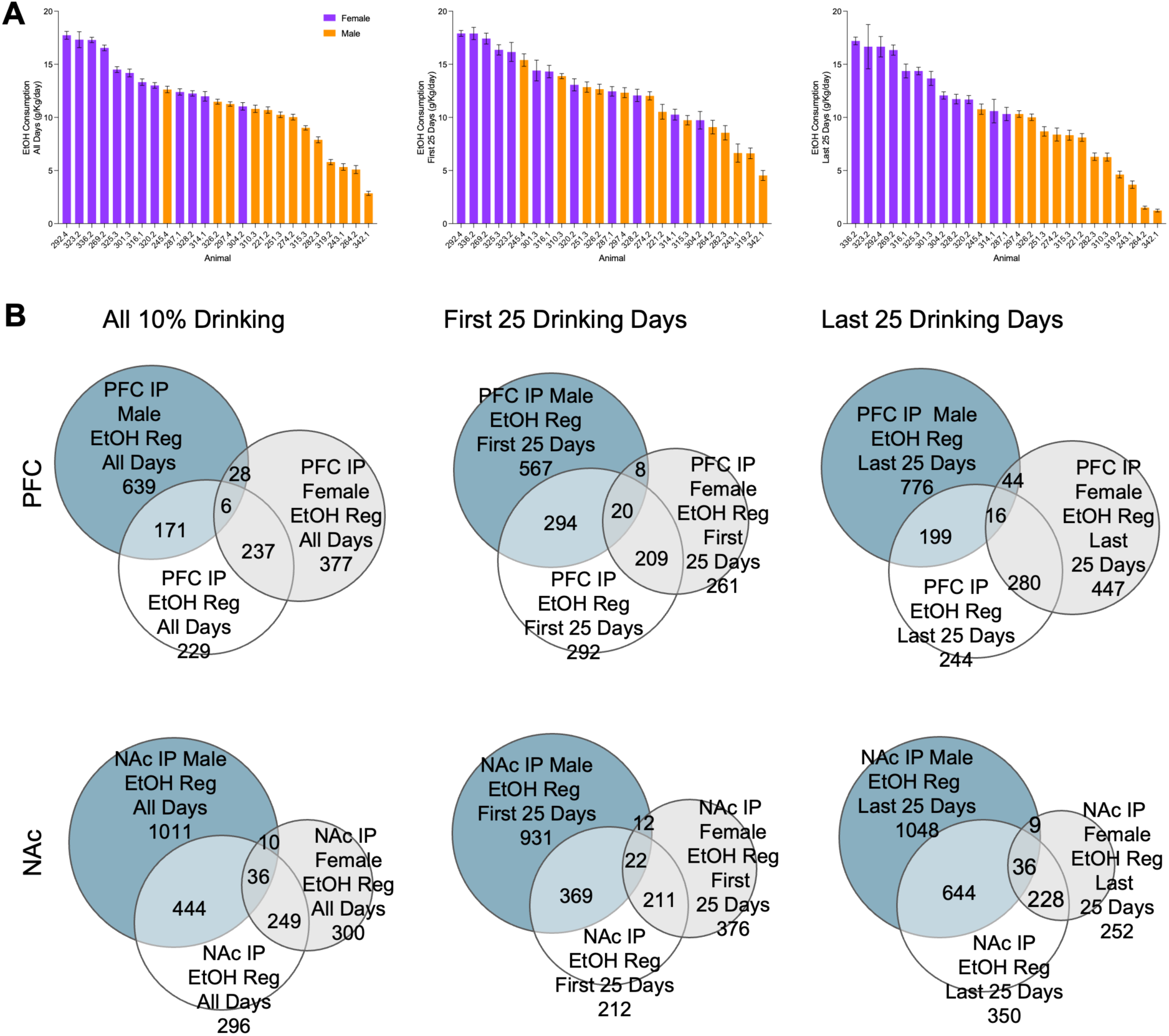
2-BC drinking for all EtOH drinking animals. **A**. Individual animal average drinking is shown across all 10% ethanol drinking days, the first 25 days, and last 25 drinking days. Females are shown in purple and males are colored orange. **B**. The relationship between drinking level and astrocyte gene translation was assessed based on all 10% EtOH drinking days, the first 25 EtOH drinking days, and the final 25 EtOH drinking days in the PFC and NAc. Overlap between DT astrocyte genes when males and females are combined, males alone, and females alone are shown for each time-period and brain region. The numbers below the text or alone represent the number of DT genes within the portion of the Venn diagram.

Individual drinking levels shown in Figure 3A were used in a gene-wise linear model to identify genes with positive or negative correlations with drinking for the PFC and NAc (Figure 3B). In the PFC, using the drinking level during the first 25 days resulted in the highest number of significantly regulated genes in the astrocyte fraction when both sexes were included in the comparison (Table 1). In the NAc, drinking during the last 25 days resulted in the highest number of regulated genes in the astrocyte fraction in the combined sexes comparison (Table 1). Overlap between males and females for both regions and drinking timeframes is minimal suggesting that chronic EtOH drinking alters astrocyte translation differently in males and females.

The top 30 enriched pathways in the PFC associated with EtOH drinking included ‘Extracellular matrix organization’, ‘Glycation Signaling Pathway’, and ‘Axonal Guidance Signaling’ (Figure 4A). In the PFC of females, calcium signaling had a negative activation z-score while potassium channels had a positive activation profile. Other pathways of note in the analysis of the PFC of females were ‘Glycerophospholipid biosynthesis’ and ‘Neurovascular Coupling Signaling Pathway’ (Figure 4B). In the PFC of males, ‘Sphingosine-1-phosphate Signaling’ and ‘WNT/SHH Axonal Guidance Signaling Pathway’ were top enriched pathways, both with negative activation z-scores. Other pathways of note were ‘Extracellular matrix organization’, cholesterol related pathways, and ‘Neuregulin Signaling’ (Figure 4C).

**Figure 4.**
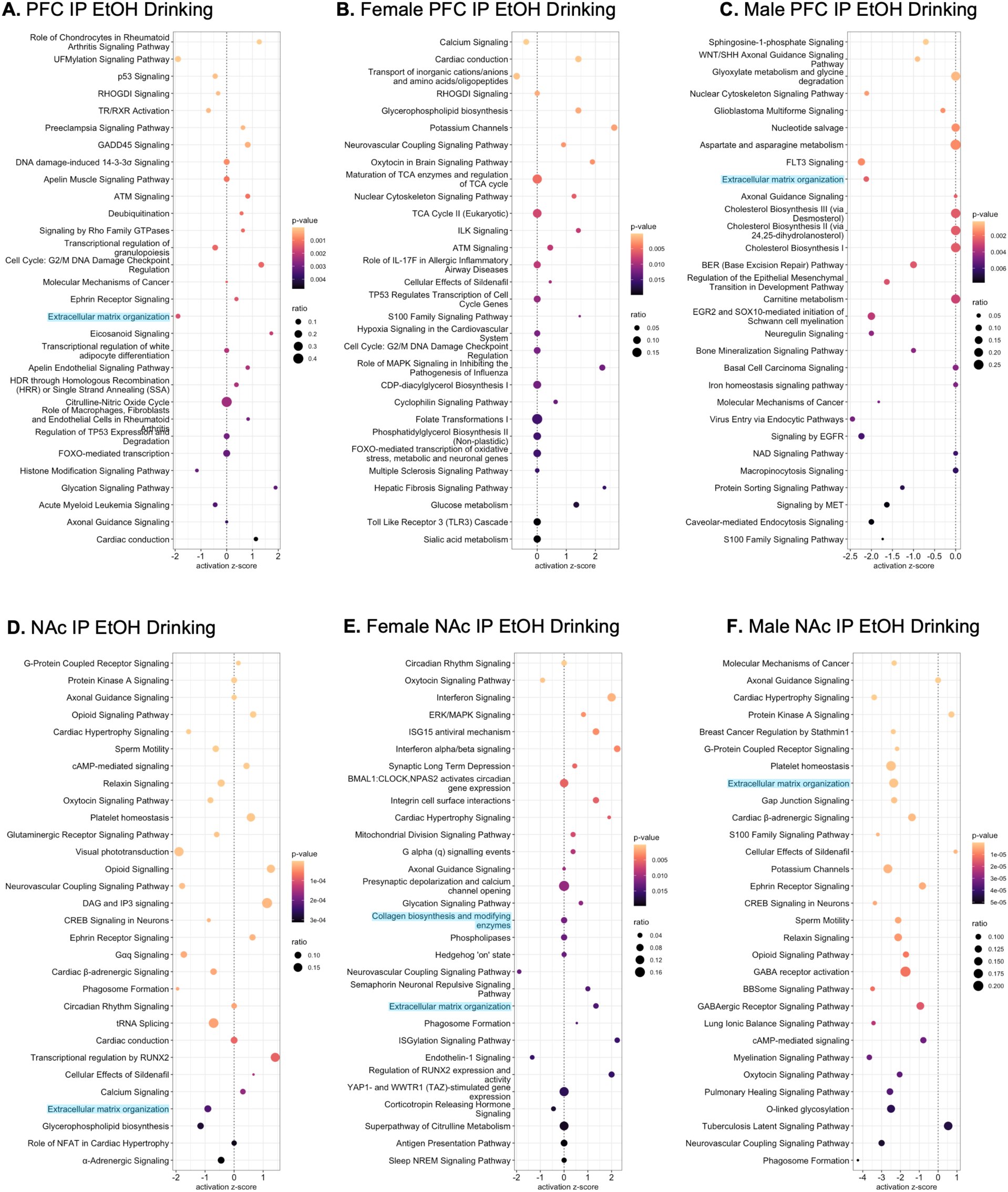
Genes with significant correlation with EtOH drinking in the astrocyte fraction across all days in astrocytes were used for IPA assessment of canonical pathway enrichment. **A**. Top 30 enriched canonical pathways in PFC associated with EtOH drinking. **B**. Top enriched canonical pathways in genes correlated with EtOH drinking in the PFC of females only. **C**. Top enriched canonical pathways in genes correlated with EtOH drinking in the PFC of males only. **D**. Top 30 enriched canonical pathways in NAc associated with EtOH drinking. **E**. Top enriched canonical pathways in genes correlated with EtOH drinking in the NAc of females only. **F**. Top enriched canonical pathways in genes correlated with EtOH drinking in the NAc of males only. Pathways related to the ECM are highlighted.

The top enriched pathway in the NAc associated with EtOH drinking was ‘G-Protein Coupled Receptor Signaling’ (Figure 4D). Other enriched pathways of note were ‘Axonal Guidance’, ‘Opioid Signaling Pathway’, and ‘Extracellular Matrix Organization’. In the NAc of females, the top enriched pathway was ‘Circadian Rhythm Signaling’ followed by ‘Oxytocin Signaling Pathway’ and ‘Interferon Signaling’. In addition, ‘Axonal Guidance Signaling’, ‘Glycation Signaling Pathway’, and ‘Extracellular matrix organization’ were enriched pathways for the NAc for females (Figure 4E). In the NAc of males, the top 30 enriched pathways included ‘Axonal Guidance Signaling’, ‘Extracellular matrix organization’, Gap Junction Signaling’, and ‘Opioid Signaling Pathway’ (Figure 4F).

### 3.5 TRAP-qPCR confirmation studies

As enriched categories related to the ECM were found to be significant in several of the IPA analyses (highlighted in Figures 2 and 4), we selected ECM-related genes that were differentially translated in our TRAP-seq analyses for confirmation by TRAP-qPCR. We confirmed that the translation of *Ncan*, encoding for neurocan, a brain-specific CSPG of the lectican family that is enriched in astrocytes, was upregulated by 2-BC drinking in the male PFC; *Csgalnact1*, encoding for Chondroitin Sulfate N-acetylgalactosaminyltransferase 1, a CS-GAG biosynthetic enzyme, and *Tnc*, encoding for tenascin C, an ECM glycoprotein that can be part of the PNNs, were down-regulated in the NAc of EtOH-drinking males; *Mmp14*, encoding for matrix metalloproteinase 14, a membrane-bound ECM proteolytic enzyme, displayed a sex-by-treatment interaction, with an increased translation in the PFC of females and a decreased translation in the PFC of males that underwent chronic drinking; *Bcan*, encoding for brevican, the other brain-specific CSPG of the lectican family also enriched in astrocytes, displayed a trend toward a sex-by-treatment interaction (Figure 5 A-D). However, the down-regulation of *Chst11*, encoding for another CS-GAG biosynthetic enzyme in the male NAc observed by TRAP-seq was not confirmed by TRAP-qPCR.

**Figure 5.**
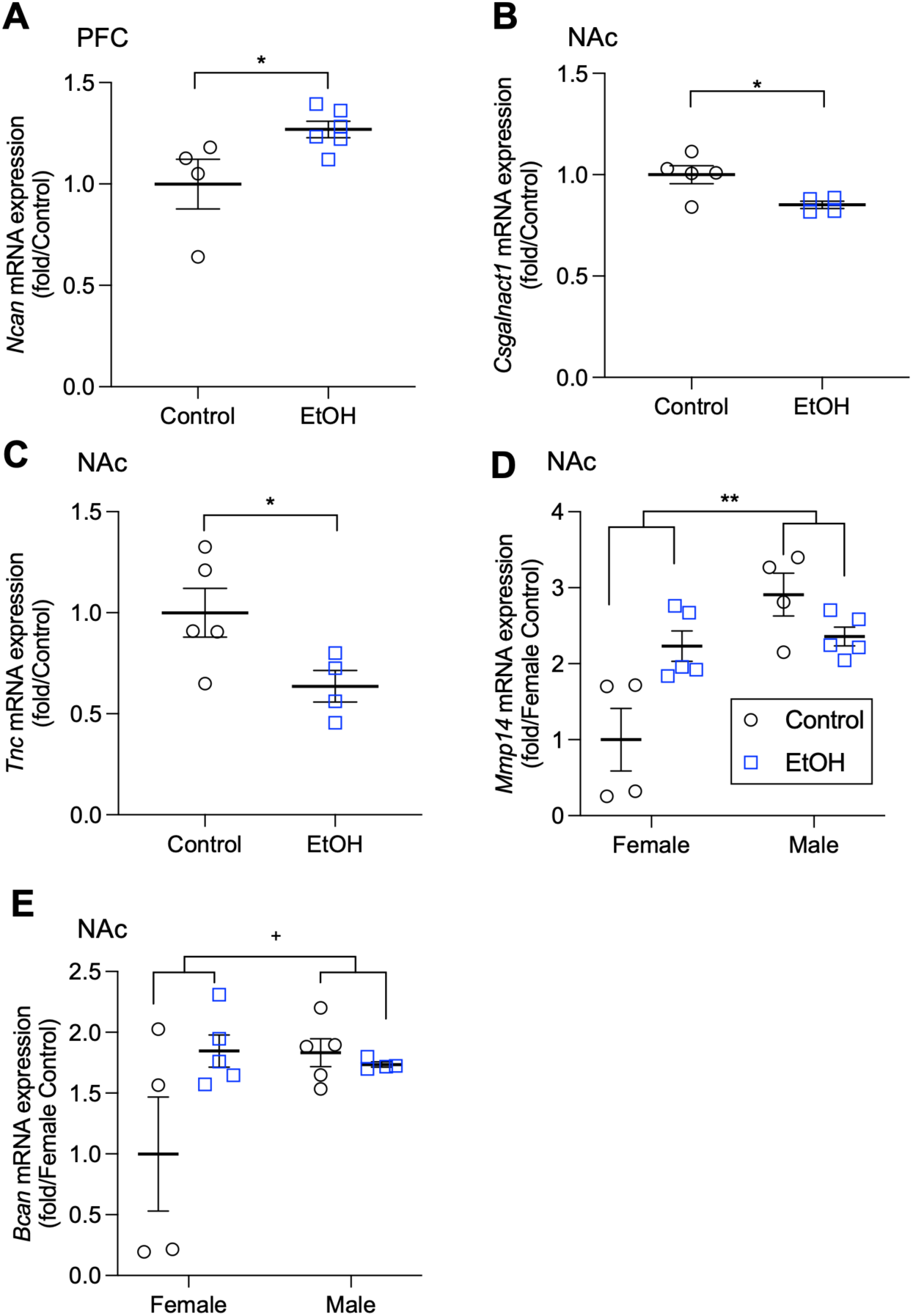
Confirmation of TRAP-Seq DT genes by TRAP-qRT-PCR. **A.** *Ncan* up-regulation by EtOH drinking in male PFC (*, p = 0.040). **B.** *Csgalnact1* down-regulation by EtOH drinking in male NAc (*, p = 0.026). **C.** *Tnc* down-regulation by EtOH drinking in male NAc (*, p = 0.049). **D.** *Mmp14* interaction of sex by treatment in PFC (**, p = 0.004). **E.** Trend for interaction of sex by treatment for *Bcan* in NAc (+, p = 0.058).

### 3.6 Glycosaminoglycans (GAGs) altered by EtOH

In our analysis of astrocyte DT by EtOH drinking vs water drinking and correlated effects with level of EtOH 2-BC drinking we observed enrichment of genes related to the ECM, axonal guidance, and neurite outgrowth. GAGs are major components of the ECM and impact axonal guidance, neurite outgrowth, and synaptic functions. For these reasons, we analyzed an additional cohort of astrocyte-TRAP mice for 2-BC drinking, using methods described in Figure 1A, to determine the effects of drinking on GAGs levels. In addition, we determined the effects of 2-BC drinking on GAGs in HP mice, which were selectively bred for high EtOH preference (a behavioral phenotype that strongly correlates with EtOH intake; Anderson et al., 2025). Similar to what we observed in the TRAP-RNA-seq study (Figure 1), astrocyte-TRAP mice drank high levels of EtOH, with females having higher overall EtOH consumption and preference (Figure 6A-B). Female HP mice had even higher EtOH consumption compared to astrocyte-TRAP mice of both sexes and HP males (Figure 6C-D). Individual animal average EtOH consumption levels are shown in Figure 6E (astrocyte-TRAP) and Figure 6F (HP).

**Figure 6.**
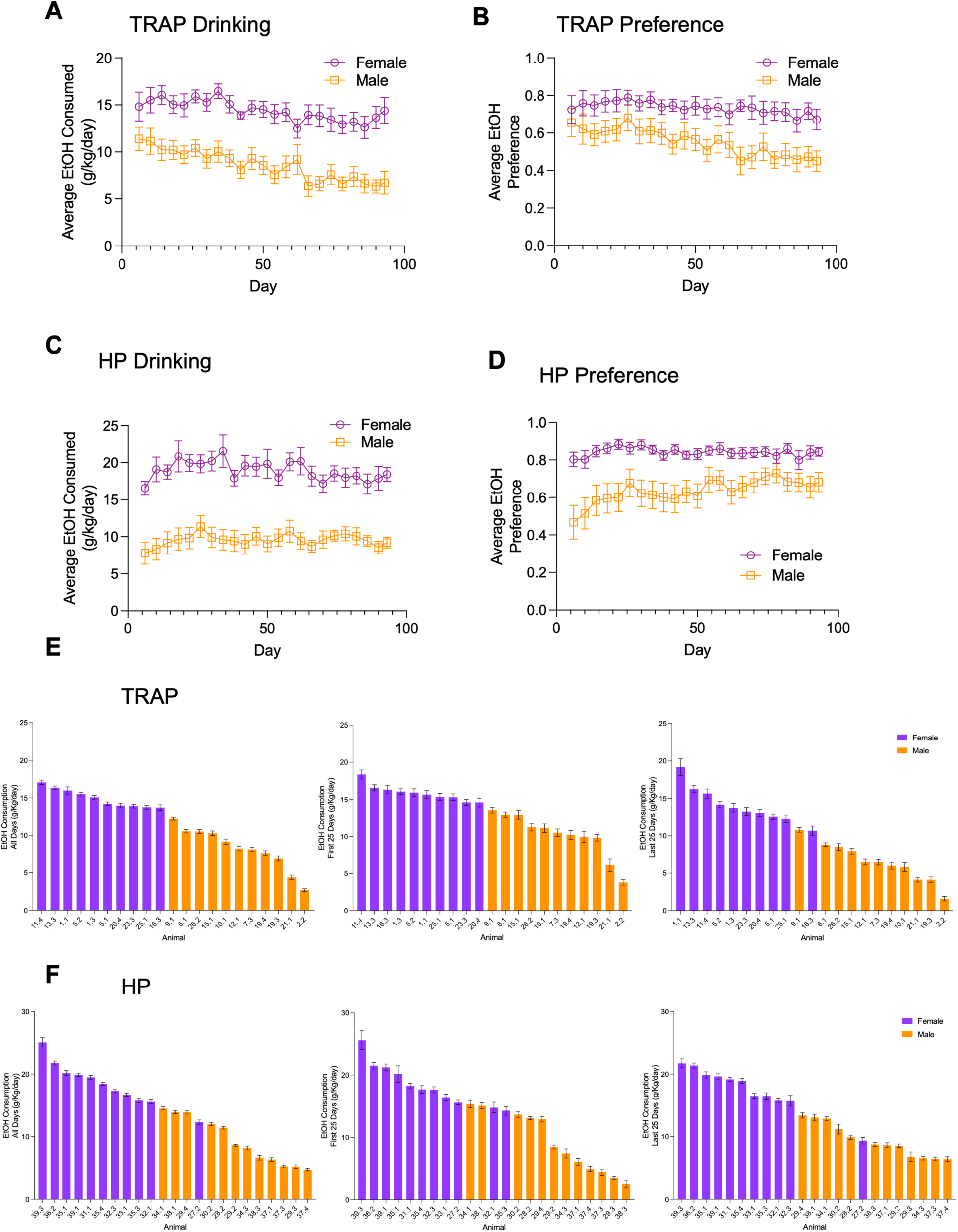
Aldh1l1-EGFP/Rpl10a (TRAP) and HP mice 2-BC drinking for GAGs analysis in NAc and PFC. In the TRAP mice average EtOH consumption. (**A**) and average EtOH preference (**B**) during the 10% EtOH drinking days in male and female TRAP is shown. Average EtOH consumption and preference ratio for HP are shown (**C**, **D**). Individual animal average drinking is shown across all 10% EtOH drinking days, the first 25 days, and last 25 drinking days for TRAP (**E**) and HP (**F**) mice.

Disaccharide analysis of CS-, HS-, and HA-GAGs revealed that in the adult NAc and PFC of male and female mice of all three lines, CS-GAGs were the most abundant GAGs, ranging between 77.96 and 79.86% of the total GAGs in the NAc and between 82.51 and 84.1% in the PFC; HS-GAGs ranged between 11.45 and 12.94% of the total GAGs in both areas and both sexes across the three lines; HA-GAGs ranged between 8.2 and 9.56% of the total GAGs in the NAc, and 3.89 and 5.45% in the PFC (Supplemental Figure 1A-C). These data suggest that GAG composition, especially CS-GAGs and HA-GAGs, varies across brain areas.

Supplemental Figure 2A-C shows the percentage distribution of differentially sulfated CS-GAGs; the most abundant CS modifications across brain regions, sexes, and lines was CS-4S (around 90% of the total CS-GAGs), which has been associated with inhibition of plasticity (Carulli et al., 2005). The other CS disaccharides identified, from the most represented to the least, were: CS-0S, CS-6S, CS-4S6S, CS-2S6S, CS-2S4S, and CS-TriS. In the PFC of astrocyte-TRAP mice, there was a significant sex by 2-BC drinking group interaction for the proportion of CS-4S, CS-0S, and CS-6S (Figure 7A). Long-term continuous access 2-BC EtOH drinking increased the proportion of NAc chondroitin sulfate sulfated at the 4S position (CS-4S) in EtOH drinking astrocyte-TRAP mice compared to water drinking controls while the proportion of CS-6S was decreased in EtOH drinking mice (Figure 7B, left and center panels). The only change in CS-GAGs observed after chronic drinking in HP mice was in the proportion of CS-TriS for which there was a significant sex by 2-BC drinking group interaction (Figure 7C, left panel); it should be pointed out, however, that this disaccharide represents only 0.2-0.4% of the total CS-GAGs.

**Figure 7.**
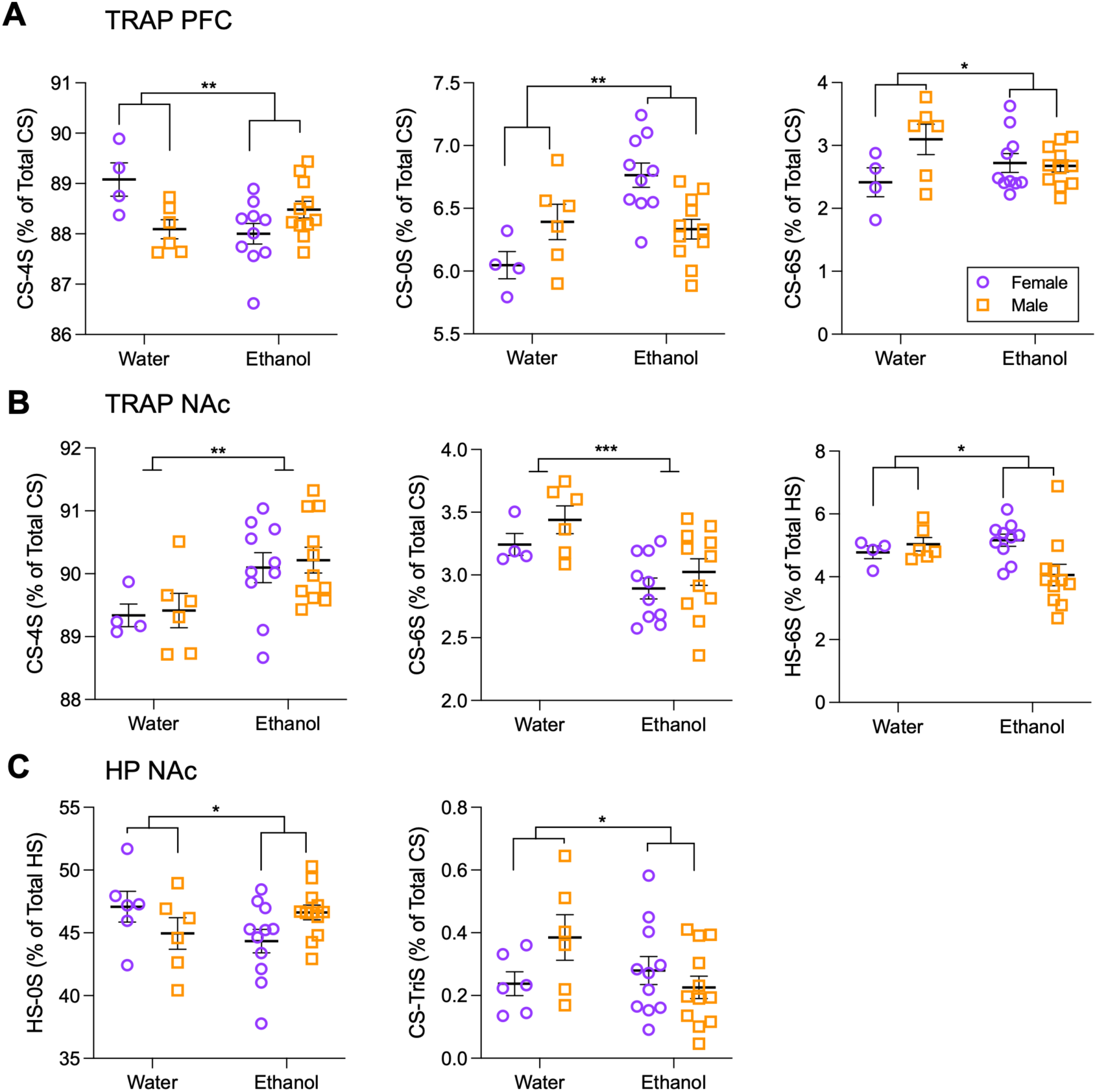
EtOH drinking alters the relative distribution of CS– and HS-GAGs in the PFC and NAc of TRAP and HP mice. **A**. In the PFC, there were sex by EtOH interactions for CS-4S, CS-0S, and CS-6S. **B**. In the NAc, EtOH resulted in higher proportion of CS-4S and lower proportion of CS-6S. For HS-6S there was an interaction of sex and EtOH drinking. **C**. In HP mice, GAGs show a sex by EtOH interaction for NAc HS-0S and CS-TriS proportion.

Supplemental Figure 3A-C shows the percentage distribution of differentially sulfated HS-GAGs. The most abundant HS modification across brain areas, sexes, and lines is HS-0S, followed by HS-NS, HS-NS-2S, HS-TriS, HS-NS6S, HS6S and HS-2S6S. Fewer changes were observed in HS-GAGs than in CS-GAGs after chronic EtOH drinking. There was a significant sex by treatment interaction for proportion of HS-6S in the NAc of astrocyte-TRAP mice (Figure 7B, right panel). In the NAc of HP mice, there was a significant sex by 2-BC drinking group interaction for proportion of HS-0S (Figure 7C, right panel).

### 3.7 GAG Differences Between HP and LP Lines

To investigate the potential role of GAG sulfation on predisposition for EtOH preference phenotypes, we compared GAGs levels in the PFC and NAc of HP and LP mice in EtOH-naïve mice. There was a significant main effect of line for CS-4S, CS-0S, CS-4S6S, CS-2S4S, and CS-TriS in the PFC. In addition, there was a significant main effect of sex for CS-6S (Figure 8A). In the NAc, there was a significant main effect of line for CS-4S, CS-0S, CS-6S, CS-4S6S, CS-2S6S, and CS-TriS proportions for data from EtOH-naïve HP and LP mice (Figure 8B).

**Figure 8.**
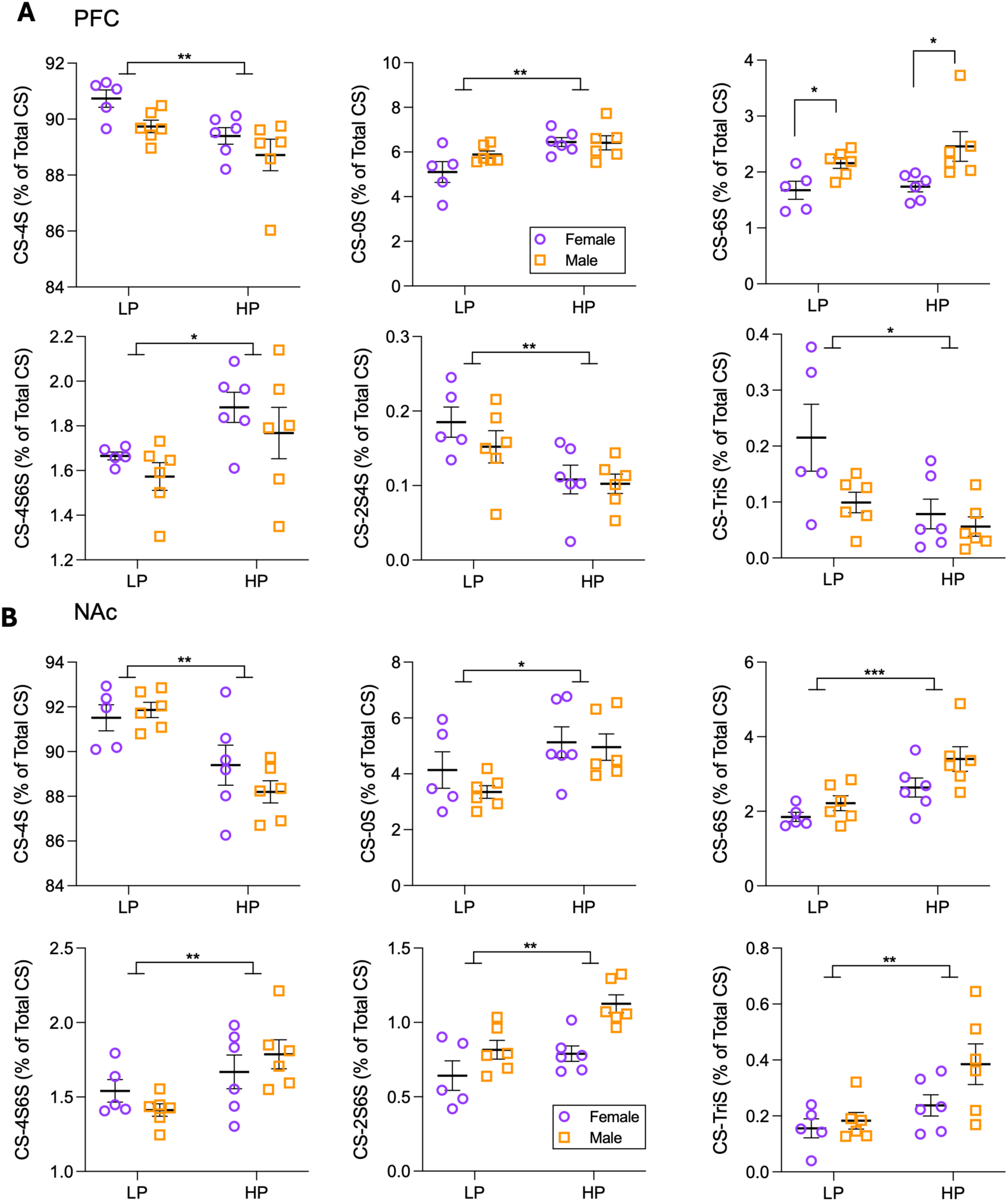
Comparison of the percentage distribution of CS-GAGs in water-drinking HP and LP mice. **A**. CS GAGs differences between HP and LP in PFC of EtOH naïve mice. Main effects of line are seen in CS-4S, CS-0S, CS-4S6S, CS-2S4S, and CS-TriS. CS-6S shows a sex difference but no effect of line. **B**. CS GAGs differences between HP and LP in NAc of EtOH naïve mice. Main effects of line are seen in CS-4S, CS-0S, CS-6S, CS-4S6S, CS-2S6S, and CS-TriS.

## Discussion

Our findings reveal that chronic EtOH consumption induces profound, sex-specific astrocytic adaptations in the PFC and NAc. Female mice consistently consumed more EtOH than males, and astrocytic translational profiles reflected this behavioral difference. Hundreds of DT genes were identified in both brain regions, with minimal overlap between astrocytic (TRAP) and bulk (input) fractions, underscoring the importance of cell type-specific analyses.

Pathway analysis highlighted significant involvement of ECM and synaptic remodeling processes. In the PFC, the complement system and glutamatergic signaling emerged as top pathways, whereas in the NAc, phagosome formation, opioid signaling, and mitochondrial function were enriched. These findings suggest astrocytic contributions to both neuroinflammatory and metabolic processes associated with AUD. Importantly, sex-specific differences were observed: females had enrichment in calcium and glutamatergic pathways, while males had alterations in axonal guidance and mitochondrial processes.

Transcriptional profiles of brain tissues from post-mortem studies (AUD vs control) have revealed that genes and biological networks associated with AUD depend on tissue and cell type (Zhang et al., 2014; Farris et al., 2015; Brenner et al., 2020; Casanova Ferrer et al., 2020; van den Oord et al., 2023). Consistent with a heterogeneous population and a complex psychiatric disorder with polygenic contributions, transcriptional meta-analyses have demonstrated tissue– and sex-specific effects, as well as substantial heterogeneity with limited overlap (Casanova Ferrer et al., 2020; Friske et al., 2025). Transcriptional profiles of brain tissues from pre-clinical animal studies have revealed that genes and biological networks associated with EtOH drinking also depend on sex, tissue, and cell type – as well as other factors (strain, species, procedural methods, etc; Casanova Ferrer et al., 2020; Hitzemann et al., 2022; Friske et al., 2025).

Even with these caveats, other transcriptomic studies have identified similar inflammatory, glutamatergic, and ECM pathways associated with risk for and consequences of chronic EtOH drinking. Colville et al. (2017, 2018) found that a former selection for high EtOH preference (as compared with low preference) performed identically to the selection for the lines used here resulted in marked effects on glutamate signaling and cell adhesion molecules (cadherins and protocadherins) in the NAc, prelimbic cortex (PL; a region within the PFC), and central amygdala (CeA). In addition, Walter et al. (2021) observed significant enrichment in genes related to translation and synaptic function, including glutamate and GABA synaptic plasticity, in the NAc core of heavy drinking rhesus macaques. Ferguson et al. (2018, 2019) found that selection for binge-like drinking in mice resulted in marked effects on inflammatory and GABAergic signaling.

Across preclinical and human studies, chronic EtOH exposure produces robust and cell-type specific transcriptional adaptations in PFC and NAc, although the precise molecular signatures vary by model and method. In the current study, chronic voluntary EtOH drinking elicited hundreds of differentially translated astrocytic genes, with stronger effects in female PFC than male, and enrichment in ECM remodeling, synaptic signaling, mitochondrial function, and immune pathways; biochemical assays confirmed EtOH-induced changes in glycosaminoglycan sulfation patterns. In a chronic intermittent EtOH vapor exposure model using male mice, Erickson et al. (2019) found that glial cells exhibited over 1150 DE genes, far exceeding changes in bulk tissue (150 DE genes). Reported findings include innate immune activation, especially type I interferon signaling, and co-expression network shifts in stress response and estrogen signaling pathways, some shared with microglia but others astrocyte-specific.

Extending to human postmortem dorsolateral PFC, single-nucleus RNA-seq carried out by Warden et al. (2025) revealed that astrocytic subtypes (Ast1–Ast3) are among the most transcriptionally dysregulated cell classes in AUD, with alterations spanning canonical astrocyte functions, neuroinflammatory signaling, and synaptic interaction programs, some of which overlapped with genetic risk loci for AUD. Together, these datasets converge on the conclusion that astrocytes in the PFC are primary responders to chronic EtOH, undergoing ECM, immune, metabolic, and synaptic gene network reorganization, with both conserved and model-specific features that may influence risk, neuroplasticity, and persistence of AUD.

In human post-mortem tissue from the NAc, van den Oord et al. (van den Oord et al., 2023) performed snRNA-Seq and identified 26 transcriptome-wide DEGs that primarily localized to medium spiny neurons, microglia, and oligodendrocytes, pointing to neuroinflammatory and myelination processes in AUD In rhesus macaque NAcc core, Walter et al. (Walter et al., 2021) reported 734 DE genes in heavy drinkers versus controls, with down-regulation of glutamatergic/GABAergic synapse terms and up-regulation of ribosomal/EIF2–mTOR signaling, consistent with broad translational control and synaptic remodeling in chronic EtOH exposure Our astrocyte-TRAP study complements and extends these observations by isolating astrocyte-specific translational changes after long-term 2-BC drinking: we observed hundreds of differentially translated genes with minimal overlap to bulk input, underscoring the value of cell type-resolved profiling. Pathway analyses identified Complement System and integrin/ECM signaling as top effects in PFC, and Phagosome Formation, opioid signaling, and mitochondrial pathways in NAc, with marked sex divergence (greater effects in female PFC, male NAc). Beyond prior work, we provide biochemical evidence that chronic drinking alters glycosaminoglycan sulfation in a region– and sex-dependent manner—implicating astrocyte-driven ECM composition as a modulator of plasticity and behavior. Finally, astrocyte translational signatures track with drinking dynamics (early PFC, late NAc), suggesting temporally evolving glial adaptations not captured in cross-sectional human or non-human primate datasets.

Our analysis of individual EtOH intake further demonstrated that transcriptomic changes correlated with drinking behavior; we found the strongest associations for PFC with early and NAc with later consumption. This finding suggests that astrocytic adaptations evolve dynamically across the course of EtOH exposure.

Lecticans (neurocan, brevican, versican, and aggrecan) consist of core-proteins attached to one or more CS-GAG chains which can be modified by sulfation and the most abundant modification in the brain is at position 4 (CS-4S) (Prydz and Dalen, 2000); (Supplemental Figure 2). CS-GAGs, and in particular CS-4S GAGs, inhibit synapse formation, axonal growth, and axon regeneration, while the digestion of CS-GAGs promotes recovery after brain and spinal cord injury (Carulli et al., 2005; Zhao and Fawcett, 2013). Comparisons of HP and LP selected lines revealed innate differences in baseline GAG composition, with HP mice having less CS-4S in both the PFC and NAc compared to LP mice, highlighting the potential role of the ECM in predisposing individuals to EtOH consumption.

We have recently found decreased CS-4S to be associated with increased neuronal complexity in neonatal mouse hippocampus after a third-trimester equivalent model of EtOH exposure (Hashimoto et al., 2025). In the current study, we found that chronic EtOH drinking altered GAG sulfation patterns in a sex– and region-dependent manner. In the PFC, we found a significant sex by treatment interaction for CS-4S, CS-0S, and CS-6S (Figure 6A) that suggests a more permissive environment for synapse formation in females (lower CS-4S in females drinking EtOH). However, the NAc of EtOH drinking mice had a significantly higher proportion of CS-4S than mice drinking water, suggesting that EtOH may decrease plasticity in the NAc by the increased production of this inhibitory GAG.

CS-GAGs are major components of the PNNs, which surround the cell body of inhibitory neurons and are associated with reduced synaptic plasticity (Carulli et al., 2005). Alteration of PNNs in response to EtOH consumption has been observed (Aguilar and Lasek, 2024). For example, six weeks of binge-like drinking increased PNNs in the insula cortex (Chen et al., 2015). PNNs have also been found to be important modulators of drinking-related behaviors and their removal by exogenous administration of chondroitinase ABC in the insula cortex decreased aversion-resistant 2-BC EtOH drinking in mice (Chen and Lasek, 2020). In addition, a recent preprint reported that chronic intermittent EtOH exposure degrades PNNs in the dorsolateral striatum and results in decreased GABAergic synapses on fast-spiking interneurons (Patton et al., 2025). The alterations caused by EtOH consumption are likely brain region-specific, and dependent on specifics of the EtOH exposure (e.g., frequency, duration).

These findings align with prior evidence implicating astrocytes in addiction (Kruyer et al., 2023; Holt and Nestler, 2024; Guizzetti et al., 2025) and extend knowledge by identifying ECM and GAG remodeling as key astrocytic adaptations to EtOH. The results underscore the importance of sex as a biological variable in AUD research. These results offer support for mechanistic investigation into the role of the ECM and excitatory/inhibitory balance in chronic EtOH drinking. Collectively, our results suggest that astrocytic adaptations in ECM and GAG pathways are critical mediators of EtOH-related neuroplasticity, providing new insights into potential therapeutic targets for AUD.

## Acknowledgements

This study was supported by NIH P60AA010760, R01AA029486, U01AA029965, U01AA013519, VA Merit Review Awards I01BX001819 and I01BX006570, Research Career Scientist Award 5IK6BX006342, and by facilities and resources at the Portland VA Health Care System. The contents of this article do not represent the views of the United States Department of Veterans Affairs or the United States government. We thank Harue Baba, Sara Aldrich, Mackenzie Jenkins, Venice Loar, Ian Anderson, Luis Tzab, David Madison, Sara Aldrich, Denesa Lockwood, and Nick Margolies for their help with the drinking studies. The authors also acknowledge the support of the Oregon National Primate Research Center Bioinformatics & Biostatistics Core, which is funded in part by NIH grant OD P51 OD011092. The conceptualization and generation of the original HP and LP selected mouse lines were by Robert Hitzemann, who also contributed to the overall study design for the long-term 2-BC drinking procedure used in these studies.

---

**Supplemental Figure 1.** CS-, HS, HA-GAGs percentage distribution in the PFC and NAc of control female and male mice across three lines. **A**: TRAP mice; **B**: HP mice; **C**: LP mice.

**Supplemental Figure 2.** Percentage distribution of CS-GAG modifications in the PFC and NAc of control female and male mice across three lines. **A**: TRAP mice; **B**: HP mice; **C**: LP mice.

**Supplemental Figure 3.** Percentage distribution of HS-GAG modifications in the PFC and NAc of control female and male mice. **A**: TRAP mice; **B**: HP mice; **C**: LP mice.

