## Supplementary figures and images for "Chronic Ethanol Drinking Alters Medial Prefrontal Cortex and Nucleus Accumbens Astrocyte Translatome and Extracellular Matrix Glycosaminoglycans"

### Supplemental Figure 1

Supplemental Figure 1A

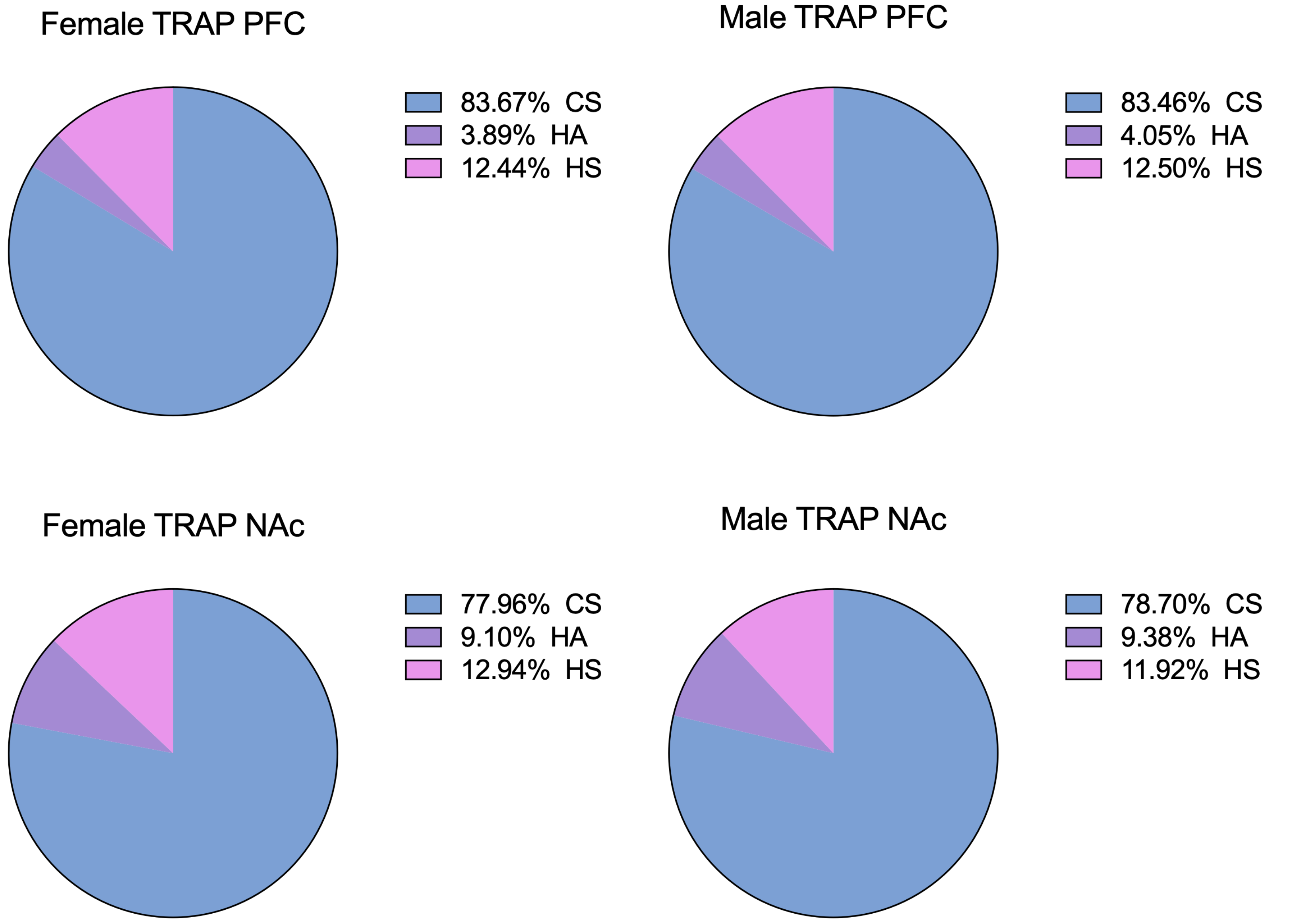

Supplemental Figure 1B

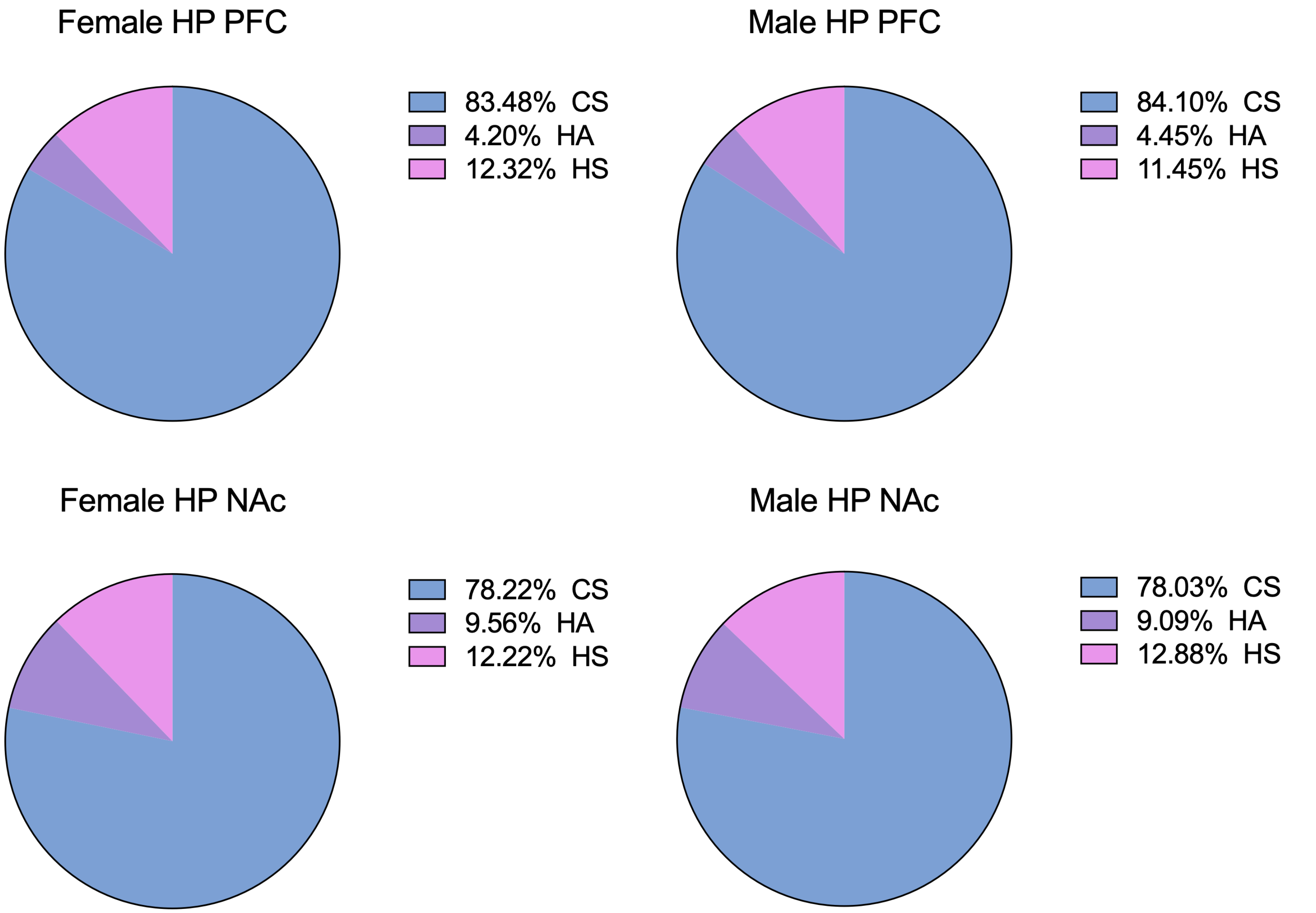

Supplemental Figure 1C

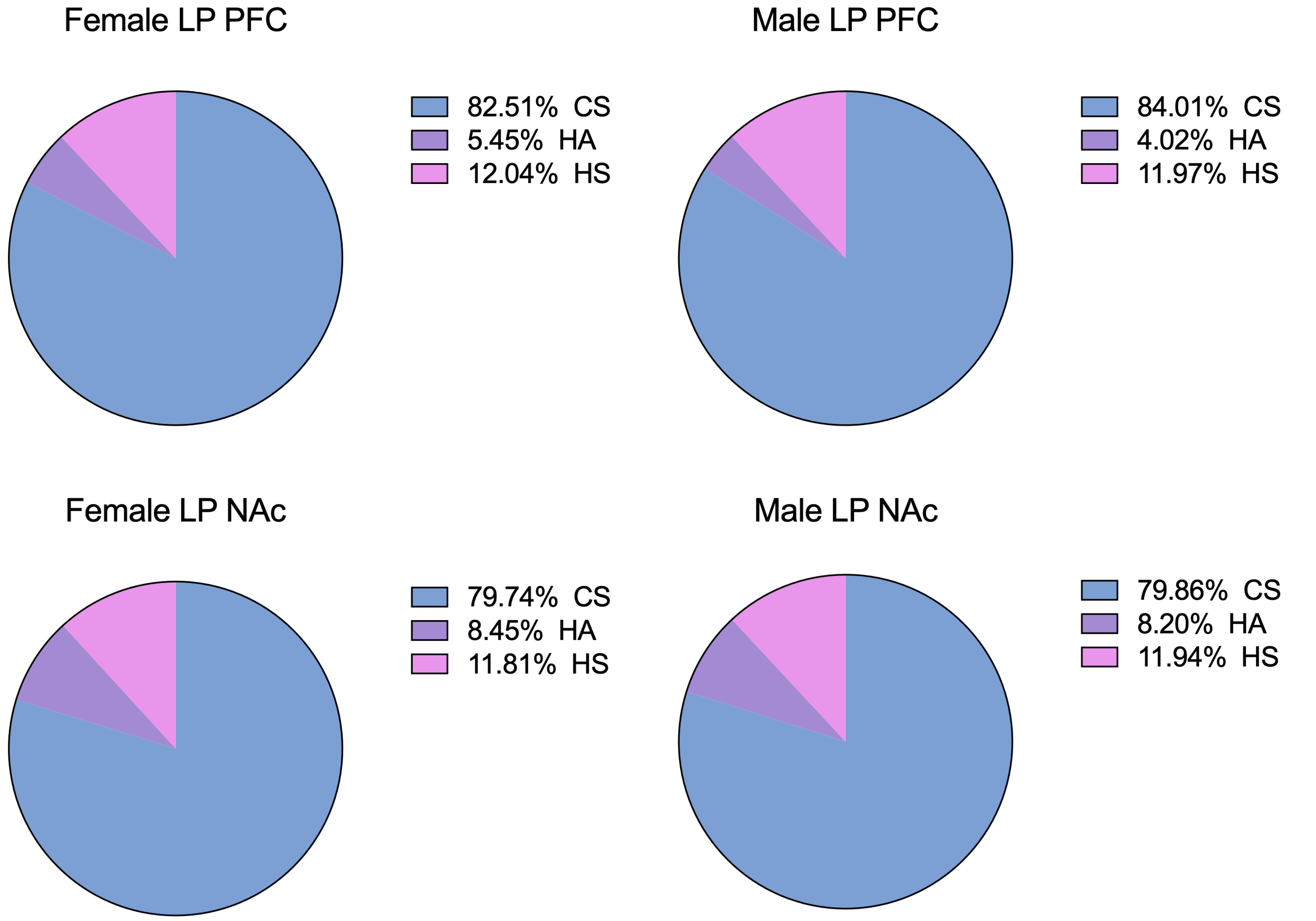

### Supplemental Figure 2

Supplemental Figure 2A

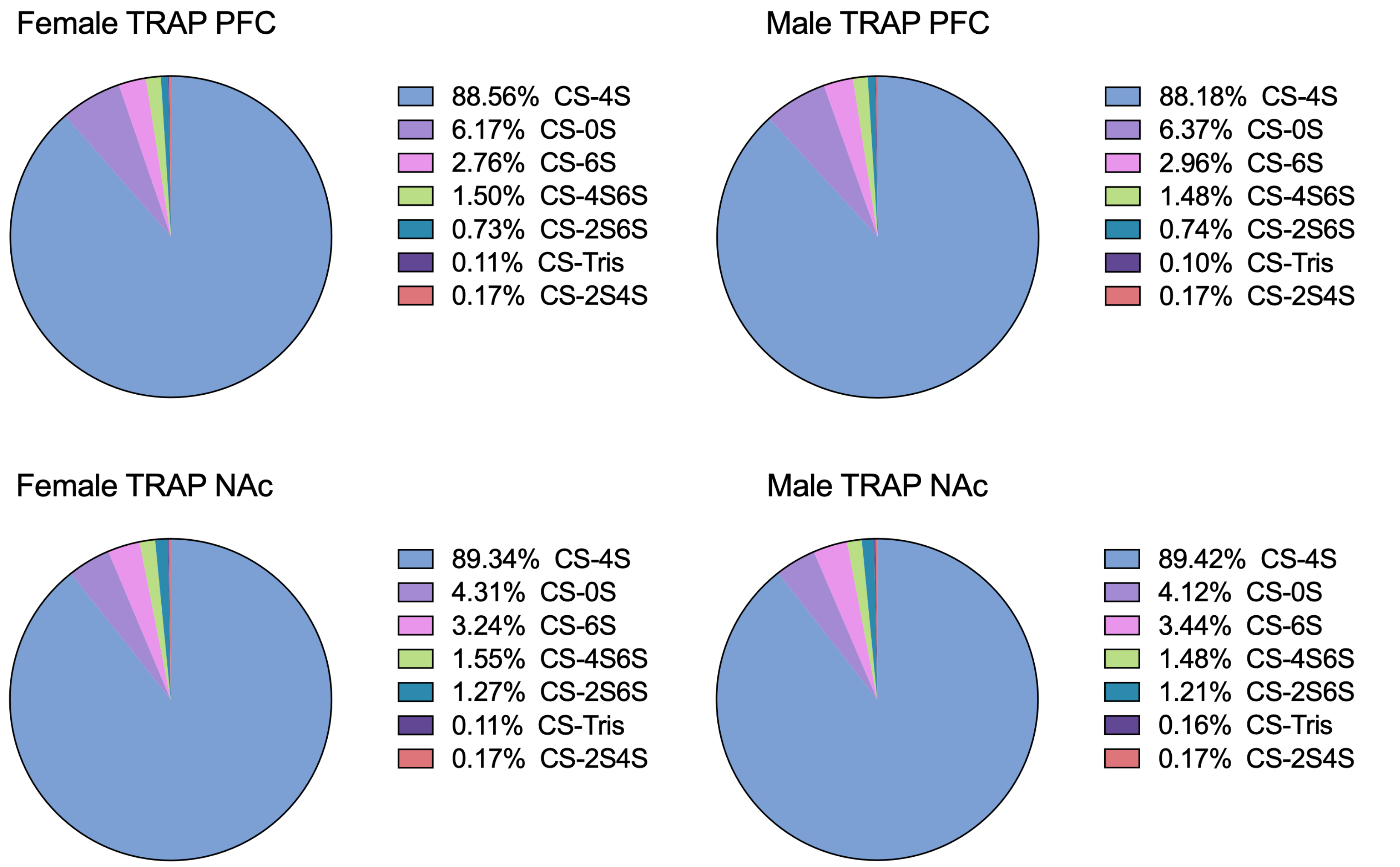

Supplemental Figure 2B

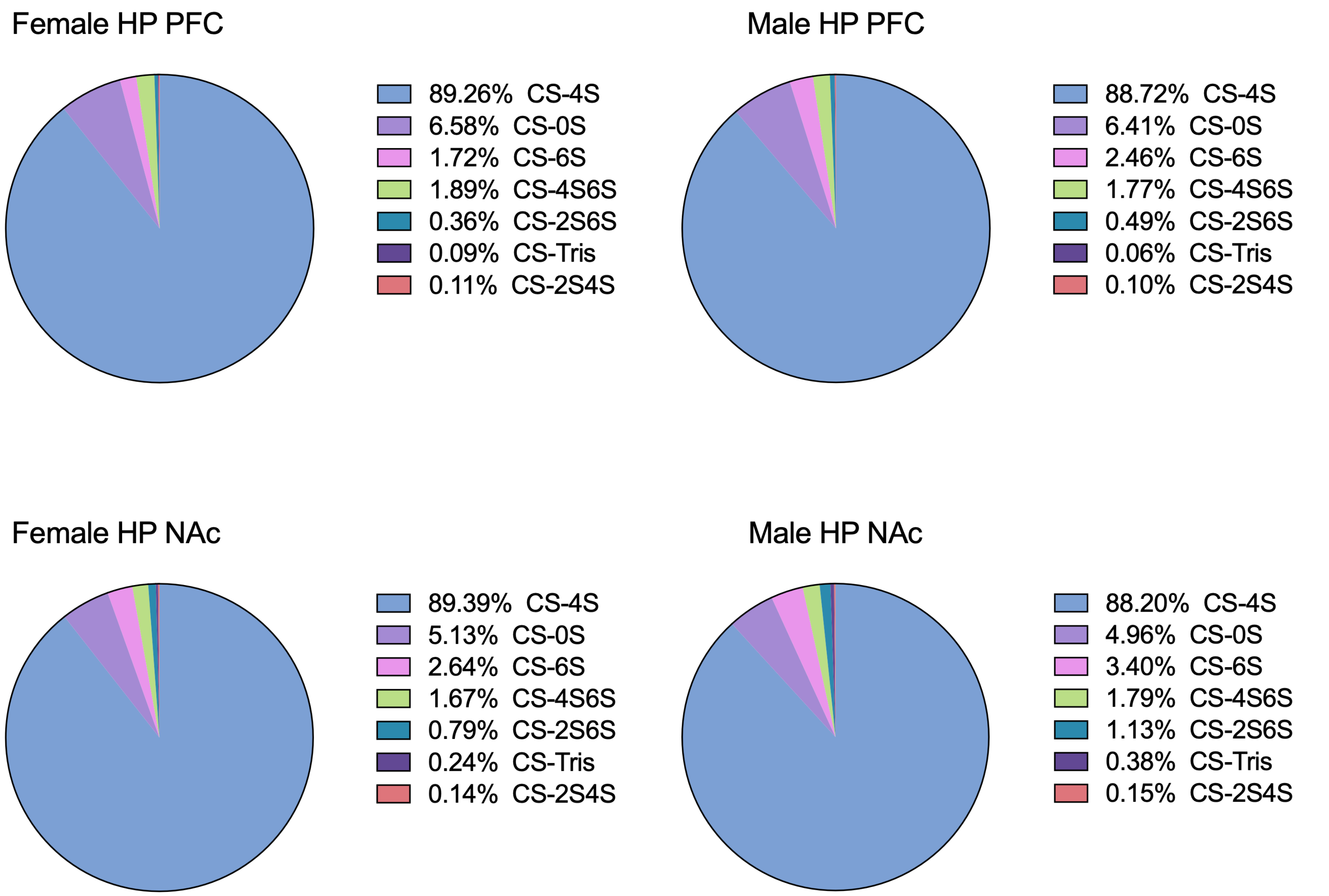

Supplemental Figure 2C

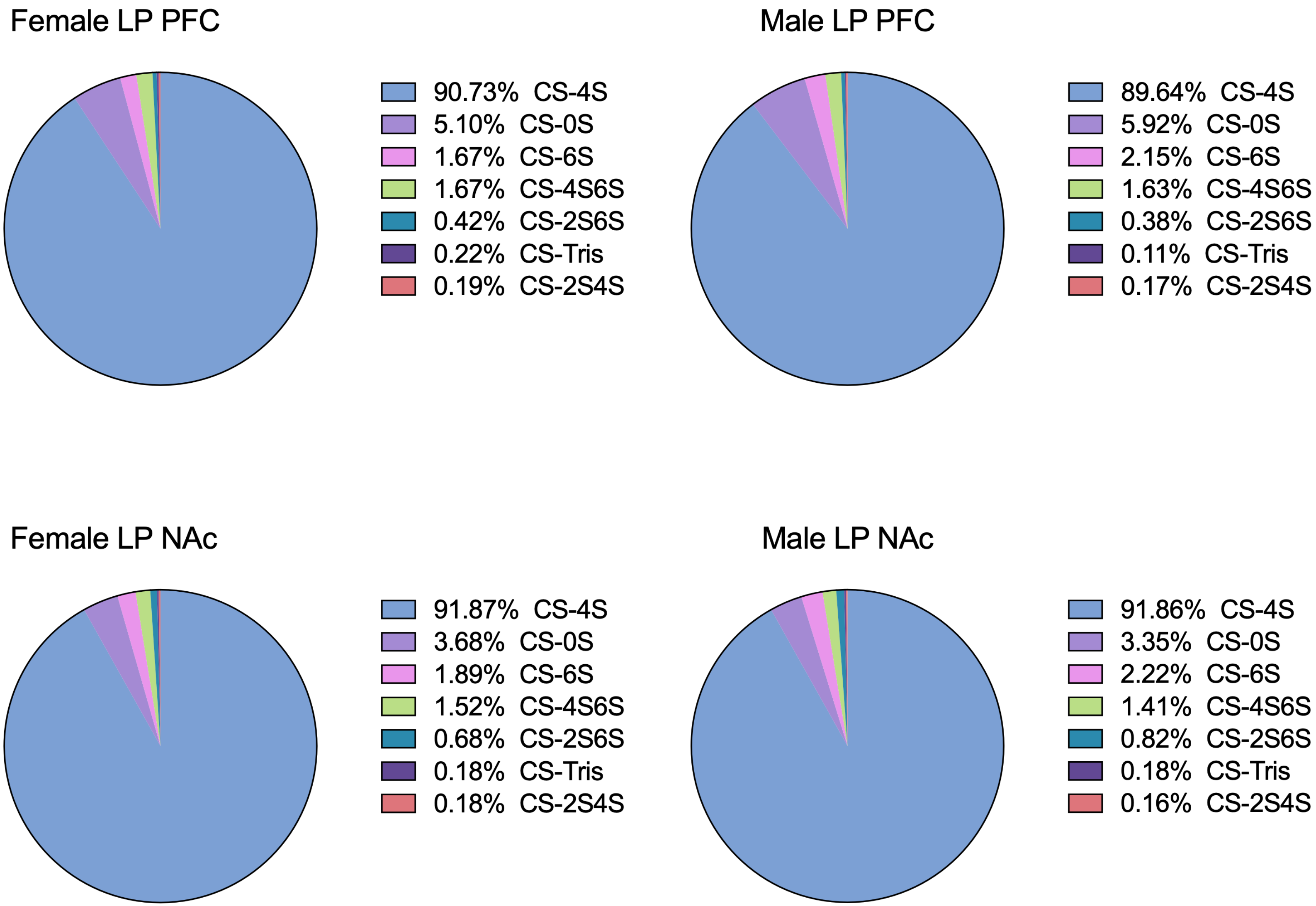

### Supplemental Figure 3

Supplemental Figure 3A

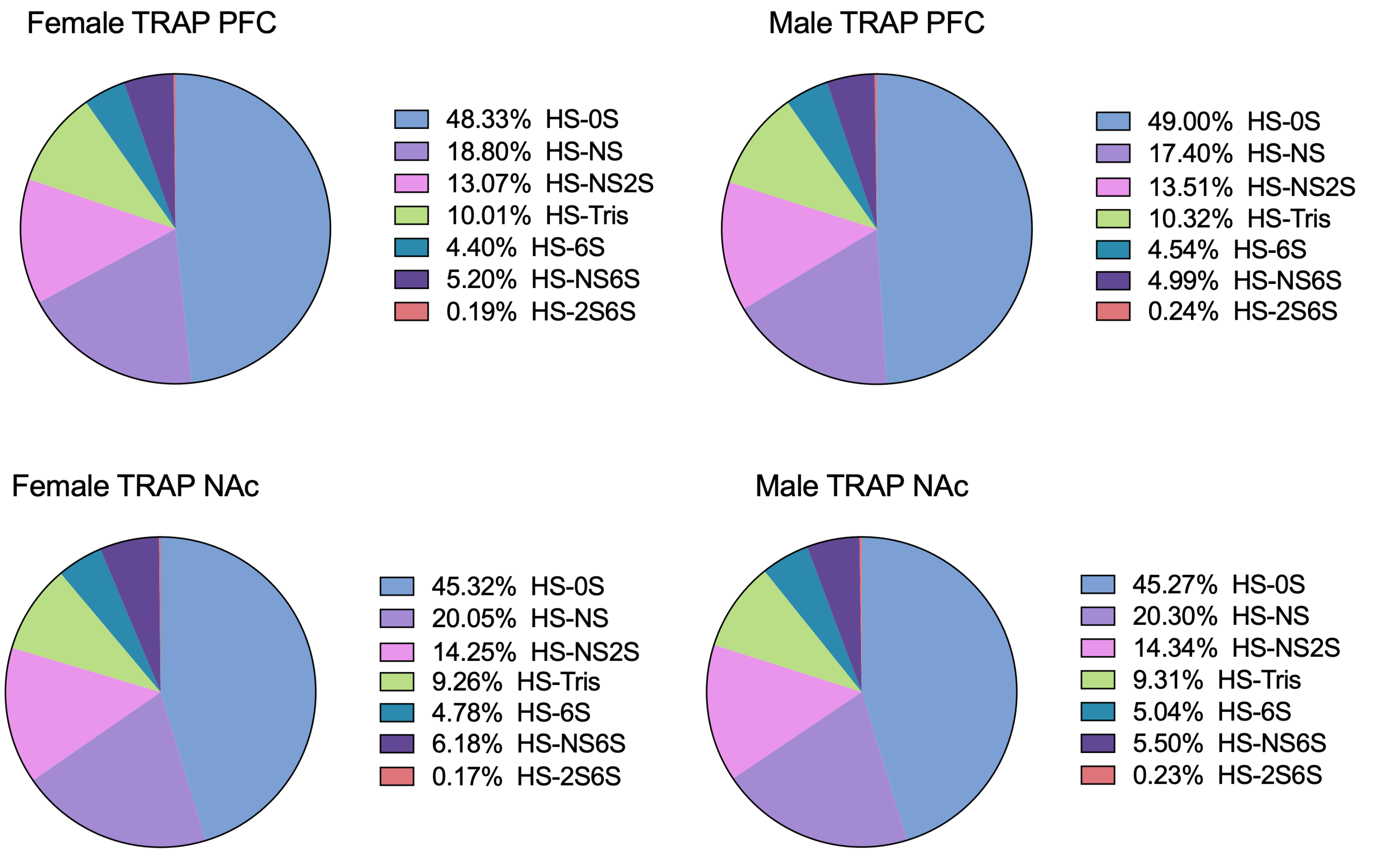

Supplemental Figure 3B

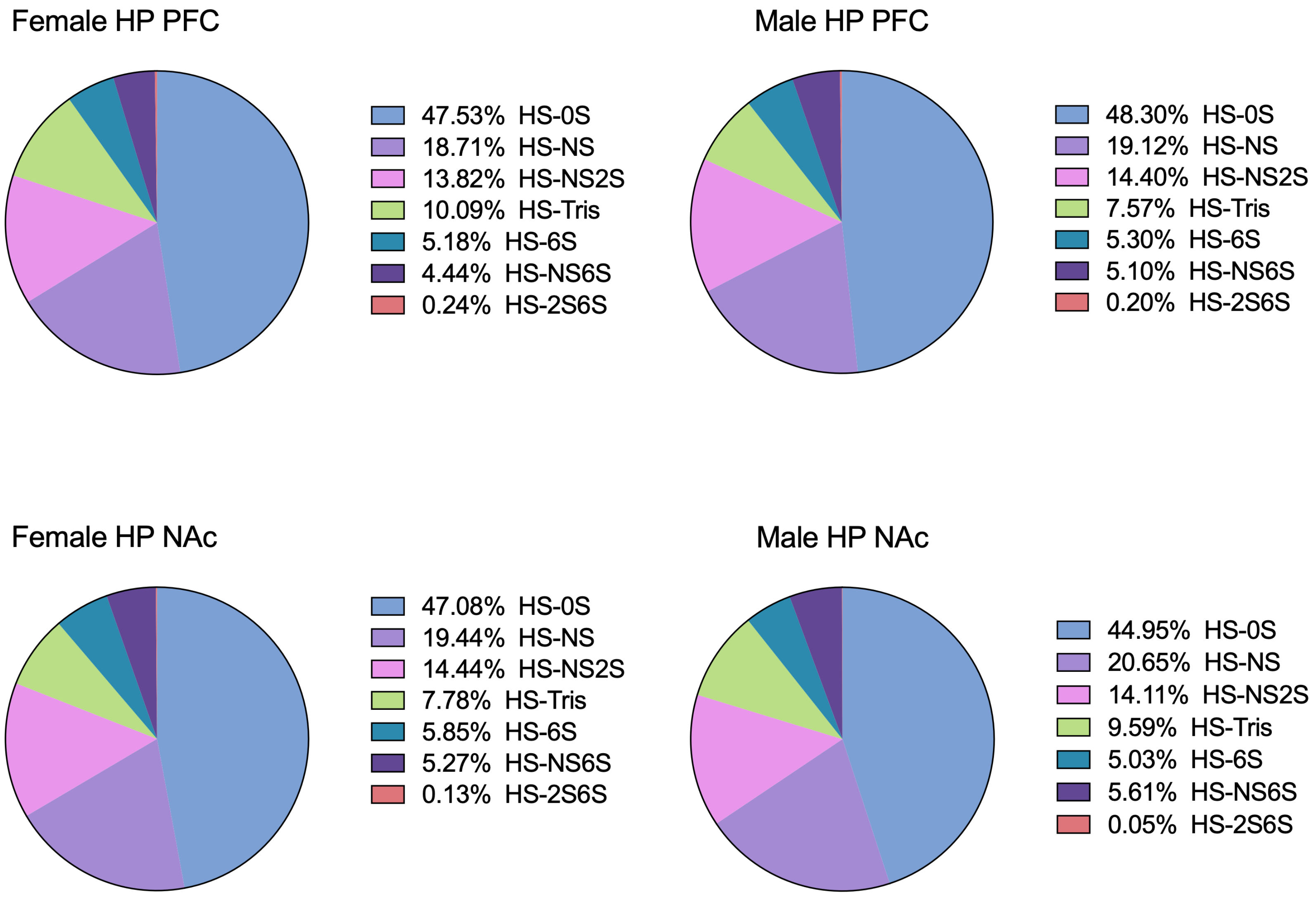

Supplemental Figure 3C

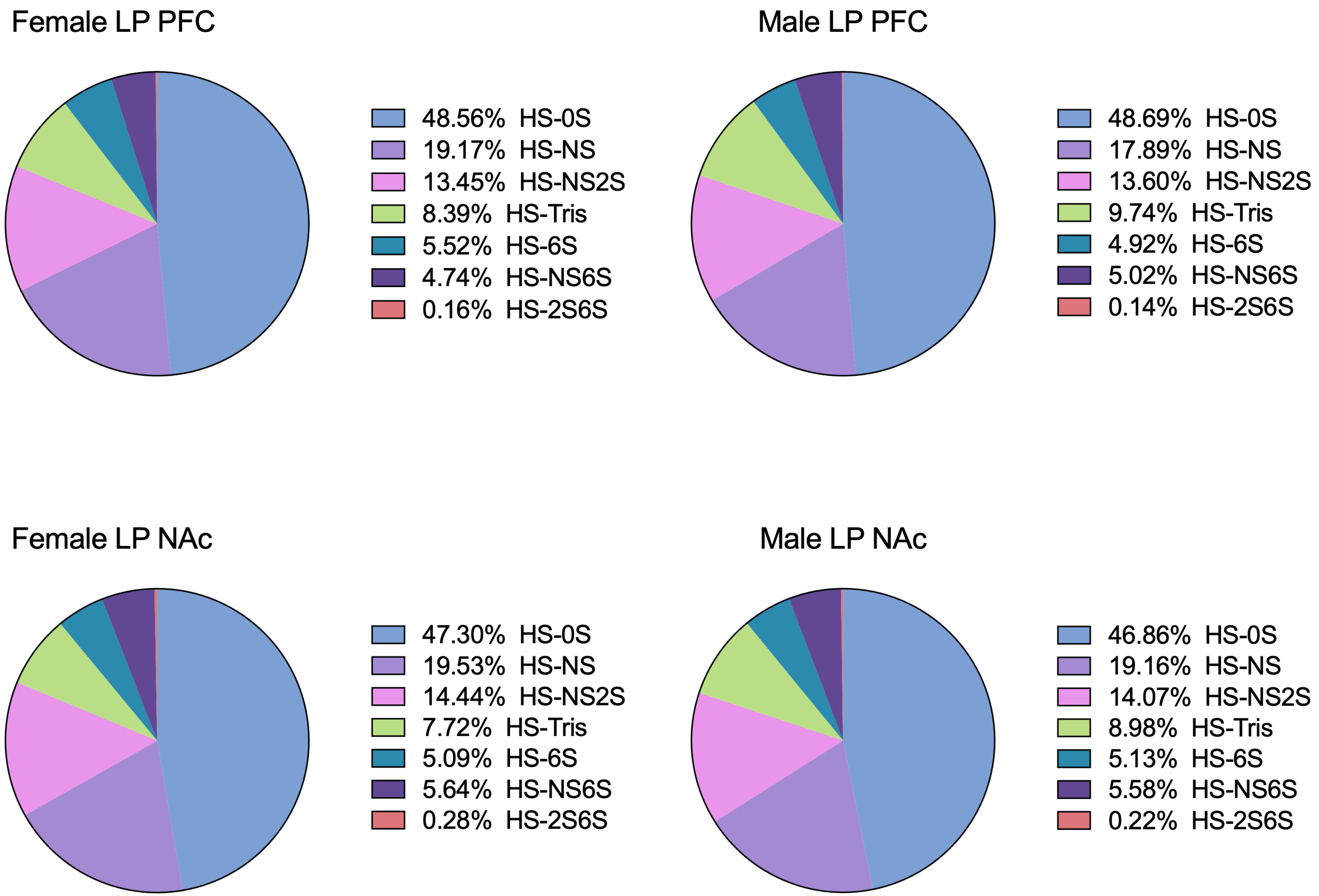
